# Combined single-cell RNA-seq profiling and enhancer editing reveals critical spatiotemporal controls over thalamic nuclei formation in the murine embryo

**DOI:** 10.1101/2022.02.18.481097

**Authors:** Kiya W. Govek, Sixing Chen, Paraskevi Sgourdou, Yao Yao, Steven Woodhouse, Tingfang Chen, Marc V. Fuccillo, Douglas J. Epstein, Pablo G. Camara

## Abstract

The thalamus is the principal information hub of the vertebrate brain, with essential roles in sensory and motor information processing, attention, and memory. The molecular mechanisms regulating the formation of thalamic nuclei are unclear. We apply longitudinal single-cell RNA-sequencing, regional abrogation of Sonic-hedgehog (Shh), and spatial profiling of gene expression to map the developmental trajectories of thalamic progenitors, intermediate progenitors, and post-mitotic neurons as they coalesce into distinct thalamic nuclei. These data reveal that the complex architecture of the thalamus is established early during embryonic brain development through the coordinated action of four cell differentiation lineages derived from Shh-dependent and independent progenitors. We systematically characterize the gene expression programs that define these lineages across time and demonstrate how their disruption upon Shh depletion causes pronounced locomotor impairment resembling infantile Parkinson’s disease. These results reveal key principles of thalamic development and provide mechanistic insights into neurodevelopmental disorders resulting from thalamic dysfunction.

## Introduction

The thalamus develops in the posterior region of the diencephalon, between the mesencephalon and telencephalon. This location is important for unique aspects of thalamic function, to process and relay sensory and motor information to and from the cerebral cortex, and to regulate sleep, alertness, and consciousness ^1^. How the thalamus comes to reside within this region of the central nervous system (CNS) has been the subject of much investigation ^2,3^. Extracellular signals secreted from key locations both extrinsic and intrinsic to the thalamic primordium have been identified and shown to play important roles in the growth, regionalization, and specification of thalamic progenitors ^4–16^. One factor in particular, the secreted morphogen Sonic hedgehog (Shh), has been implicated in spatiotemporal and threshold models of thalamic development that differ from other areas of the CNS. This impact of Shh is due, in large part, to its expression within two signaling centers, the basal plate and the zona limitans intrathalamica (ZLI), a dorsally projecting spike that separates the thalamus from the prethalamic territory ^17,18^. Shh signaling from these dual sources exhibits both unique and overlapping functions in the control of thalamic progenitor identity and specification of thalamic nuclei ^11–13^.

The adult thalamus is subdivided into 44 distinct anatomic nuclei that are categorized on the basis of their positioning (anterior, medial, lateral, ventral, posterior, and intralaminar groups), cytoarchitectural properties, types of input (sensory, motor, or limbic), and hierarchical connectivity patterns to and from the cerebral cortex and subcortical areas ^1,19–25^. Principal sensory nuclei have strong reciprocal connections with the cerebral cortex. For instance, the ventroposterior medial (VPM), dorsolateral geniculate (DLG), and ventromedial geniculate (vMG) constitute ‘first order’ thalamic nuclei that relay sensory information from the periphery along thalamocortical axons to cortical neurons within primary somatosensory, visual and auditory cortices, respectively. In turn, corticothalamic axons project to corresponding ‘higher order’ thalamic nuclei, including posterior (Po), lateral-posterior (LP) and dorsomedial geniculate (dMG), respectively. Reciprocal projections from these higher order thalamic nuclei then connect with neurons of the corresponding secondary cortical areas ^23–25^. The functions of these higher order relays are to modulate thalamocortical transmission and transfer information between cortical areas ^26^. With respect to the motor thalamus, the ventral-anterior (VA) and ventral-lateral (VL) thalamic nuclei operate as a super- integrator of driver inputs from the motor cortex, basal ganglia and cerebellum that are transmitted to cortical areas for preparation and execution of movements ^27^.

The spatial arrangement of thalamic nuclei is important for generating the precise topographical relationship needed to fulfill its role as a relay and information processing center. Despite advances in our understanding of the early events regulating thalamic growth and regionalization, there remain major gaps in knowledge of the mechanisms by which heterogeneous clusters of mostly excitatory relay neurons are specified and aggregate into distinct thalamic nuclei. One particular challenge has been to decipher the full complement of thalamic progenitor identities and to elucidate their contribution to specific thalamic nuclei ^11,12,28–32^. The results of in vivo clonal analyses indicate that individual thalamic progenitors contribute to more than one thalamic nucleus ^33,34^. The principal sensory nuclei (VPM/VPL, DLG, and vMG) and possibly the motor thalamus (VA/VL), share a similar ontology. This ontological pathway contrasts with other thalamic nuclei that segregate into three distinct clusters related to the position of thalamic progenitors during development. These findings suggest that thalamic nuclei developing in close proximity to one another are more likely to share a common lineage.

Thus far, only two distinct thalamic progenitor domains have been defined by gene expression and fate mapping studies. The caudal population of thalamic progenitors, cTh.Pro, gives rise to all glutamatergic thalamic nuclei that extend axonal projections to the neocortex ^30^. The rostral population of thalamic progenitors, rTh.Pro, comprises a narrow band of cells sandwiched between cTh.Pro and the ZLI. Thalamic neurons derived from rTh.Pro are GABAergic and contribute to the ventrolateral geniculate nucleus (VLG) and the intergeniculate leaflet (IGL), neither of which project axons to the cortex ^1,12,30,35–37^.

Multiple studies have shown that thalamic nuclei exhibit extensive heterogeneity at the level of gene expression ^2,25,34,38–41^. Nevertheless, we still lack a clear understanding of the molecular logic and developmental trajectories by which thalamic nuclei acquire their distinct identities. Here, we make use of highly parallelized single-cell RNA-sequencing and spatial profiling of gene expression to molecularly and anatomically characterize thalamic progenitor subtypes, intermediate progenitors, and post-mitotic neurons across multiple stages of mouse embryonic development. Our approach overcomes limitations of conventional single-cell transcriptomic atlases, which often lack mechanistic detail, by investigating how regional abrogation of Shh expression alters thalamic lineage progression at single-cell scale. Our findings unify models of thalamic development and provide a detailed understanding of a neurodevelopmental disorder resulting from alterations in thalamic architecture.

## Results

### Deletion of SBE1 and SBE5 abrogates *Shh* expression in the ZLI

*Shh* expression in the ZLI and basal plate of the caudal diencephalon is dependent on two Shh brain enhancers, SBE1 and SBE5 ^42^. Mouse embryos homozygous for targeted deletions of SBE1 and SBE5 (*Shh^ΔSBE1ΔSBE5/ΔSBE1ΔSBE5^*, herein referred to as *ΔSBE1/5*) fail to activate *Shh* transcription in the ZLI and basal plate after E10.0, compared to control littermates (*Shh^ΔSBE1ΔSBE5/+^*) (Fig. 1A) ^42^. Consequently, Shh signaling activity in thalamic and prethalamic territories is compromised in *ΔSBE1/5* embryos, as indicated by the loss of *Gli1* expression (Fig. 1A). Despite the absence of *Shh* expression and Shh signaling activity, a GFP reporter transgene driven by SBE1 continued to be expressed in the ZLI of *ΔSBE1/5* embryos (Fig. 1A). Moreover, genes coding for transcription factors that are normally expressed in the ZLI, such as *Pitx2* and *Foxa1*, maintained much of their expression in *ΔSBE1/5* mutants, albeit in a partially reduced area (Supplementary Fig. 1). These results indicate that the cellular integrity of the ZLI remains intact in *ΔSBE1/5* embryos and highlight the utility of this mouse model for studying the implications of Shh signaling in mammalian thalamic development.

**Figure 1.**
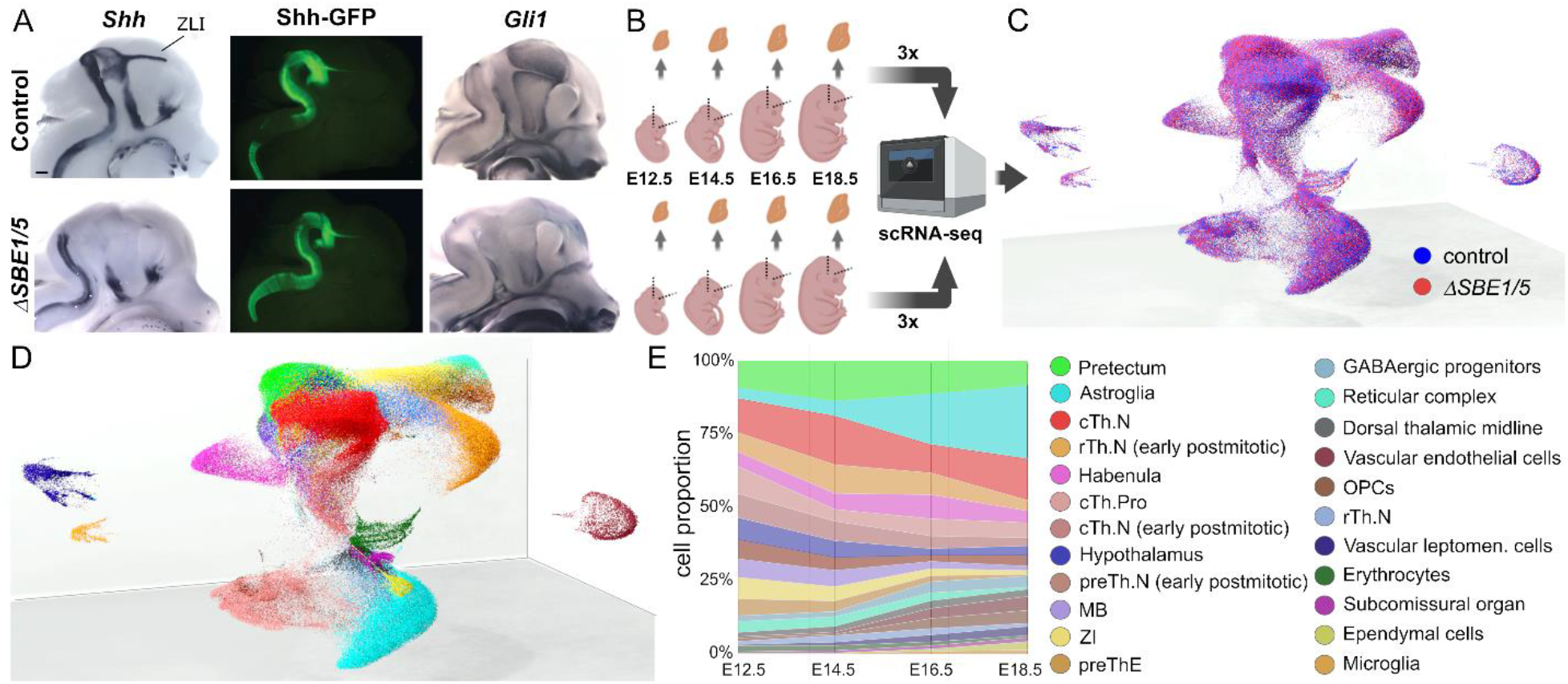
A high-resolution single-cell transcriptomic atlas of the developing caudal diencephalon. **A)** Deletion of the SBE1 and SBE5 enhancers leads to a loss of *Shh* expression and Shh signaling activity in the caudal diencephalon. Whole-mount RNA in situ hybridization for *Shh* (left) and the Shh-responsive gene *Gli1* (right) in bisected heads from control (top) and *ΔSBE1/5* embryos (bottom) at E12.5 shows reduced expression in the caudal diencephalon of mutant embryos (n=3). Scale bar = 500 μm. A Shh BAC transgene (Shh-GFP) expressing eGFP in replace of Shh shows persistent reporter activity in the ZLI of *ΔSBE1/5* and control embryos (center). **B)** Schematic of the experimental study design. The caudal diencephalon of control and *ΔSBE1/5* embryos was micro-dissected at E12.5, E14.5, E16.5, and E18.5 in three replicates per time point and genotype, fixed in methanol, and profiled with single-cell RNA-seq. **C)** 3D UMAP representation of the consolidated single-cell gene expression space across all samples, colored by the genotype of the cells. Cells from control and *ΔSBE1/5* embryos substantially overlap in the representation. **D)** 3D UMAP representation of the single-cell transcriptomic atlas colored by the 23 cell populations identified in the clustering analyses. **E)** Fraction of cells in each cell population for each of the four time points. Only cells from control mice were considered in this analysis. The observed expansion of glial cells starting between E14.5 and E16.5 is consistent with the transition between neurogenesis and gliogenesis at this stage of embryonic development. Abbreviations: cTh.N, caudal thalamic neurons; rTh.N, rostral thalamic neurons; cTh.Pro, caudal thalamic progenitors; preTh.N, prethalamic neurons; MB, midbrain; ZI, zona incerta; preThE, prethalamic eminence; OPCs, oligodendrocyte precursor cells.

### A high-resolution single-cell transcriptomic atlas of the developing caudal diencephalon in control and *ΔSBE1/5* embryos

To uncover the molecular logic driving the specification of distinct thalamic nuclei, we generated a high-resolution single-cell RNA-seq atlas of the developing caudal diencephalon in control and *ΔSBE1/5* embryos. The thalamic primordia were micro-dissected from three embryos per genotype according to anatomical landmarks (Methods) at four developmental stages (E12.5, E14.5, E16.5, and E18.5) coinciding with peak periods of thalamic proliferation, neurogenesis, and differentiation (Fig. 1B; *n* = 24 embryos in total). Overall, we captured the transcriptome of 249,076 cells (121,881 cells from control and 127,195 cells from *ΔSBE1/5* embryos) with an average of 2,597 genes detected per cell. We consolidated the single-cell gene expression space of all the samples and clustered the cells in this space (Figs. 1C and D).

Our analysis identified 23 distinct cell populations, comprising most of the known cell types in the caudal diencephalon (Figs. 1D and E). Cells from three independent biological replicates overlapped in the consolidated representation and were similarly delineated across all cell populations (Supplementary Fig. 2), indicating the absence of large batch effects. Most of the identified cell populations were part of continuous developmental trajectories, such as the differentiation of thalamic progenitors (Th.Pro) into rostral GABAergic and caudal glutamatergic thalamic neurons (denoted as rTh.N and cTh.N, respectively), or the transition from neurogenesis into gliogenesis at E14.5-16.5 (Figs. 1D and E, and Supplementary Fig. 3).

Of note, we found that these data correctly recapitulated the effect of *SBE1/5* deletions on the expression of *Shh* and Shh target genes. Expression of the basic helix-loop-helix (bHLH) transcription factor *Olig3,* distinguishes thalamic progenitors from other diencephalic cell types ^30^. As expected, differential gene expression analysis in the *Olig3*^+^ progenitor cell subpopulation showed a downregulation of Shh responsive genes (*Gli1*, *Ptch1*, *Nkx2-2* and *Olig2*) in *ΔSBE1/5* compared to control cells (Supplementary Fig. 4 and Supplementary Table 1). In addition, a detailed examination of the larger cluster of progenitor cells identified a subgroup with an expression profile consistent with the ZLI (*Shh, Foxa1, Pitx2*, *Sim2*) ^28,43^ (Supplementary Fig. 5). Differential expression analysis confirmed the strong depletion of *Shh* expression from this subgroup in *ΔSBE1/5* compared to control cells (Supplementary Fig. 5 and Supplementary Table 2). The persistence of other ZLI markers in mutant cells indicates that the ZLI is not dependent on Shh for the bulk of its formation after E10.0. Altogether, these data represent a unique resource for studies of the developing caudal diencephalon and a substantial increase in cell type resolution with respect to previous single-cell RNA-seq datasets of this brain region ^40,41,44–46^.

### Most thalamic nuclei have well-defined molecular identities at E18.5

Gene expression signatures of distinct thalamic nuclei have been described in adult mice or at single stages of embryonic development, but not in a coordinated manner across developmental time ^2,25,34,40,41,45^. We sought to systematically characterize the cell populations that define thalamic nuclei throughout embryonic development. For this purpose, we performed a separate clustering and differential gene expression analysis of E18.5 control cells from the cTh.N, rTh.N, zona incerta (ZI), habenula, and reticular complex (RT) post-mitotic cell populations. This analysis identified 23 cell subpopulations with distinct transcriptomic profiles (Figs. 2A, B and Supplementary Table 3). By comparing differentially expressed genes across subpopulations with RNA in situ hybridization data from the Allen Developing Mouse Brain Atlas ^47^, we were able to assign one or more thalamic nuclei to each of these subpopulations (Fig. 2A). Most nuclei in the thalamic, habenular, and reticular complexes were localized to distinct regions of the UMAP representation (Fig. 2A). However, in some cases, closely related thalamic nuclei, such as the anterodorsal (AD), anteroventral (AV), and anteromedial (AM), were assigned to the same transcriptomic cell subpopulation. We used a spectral graph method ^48^ to dissect the transcriptional heterogeneity within the cell populations identified in our clustering analysis and further resolved the transcriptomic signatures of some of these closely related thalamic nuclei (Fig. 2A and Supplementary Fig. 6). We were able to distinguish the transcriptomic profile of the AD/AV thalamic nuclei from that of the AM nucleus, and the profile of the VA and VL nuclei from that of the ventral medial (VM) nucleus. In other cases, such as for the ventral posterolateral (VPL) and the medial geniculate (MG) thalamic nuclei, we observed multiple transcriptomic subpopulations associated with the same nucleus, suggesting the existence of several cell populations within these nuclei. For example, we identified three distinct transcriptomic populations contributing to the MG nucleus. All of these populations expressed *Gbx2*, *Lhx2*, *Rorα*, and *Rorβ*. However, two of them had high expression of vMG markers (*Slc6a4*, *Adarb1*, *Tshz1*, *Dlk1*), whereas the other one expressed a dMG marker (*Prox1)* ^25^. Taken together, these data unveil unique molecular signatures that distinguish most thalamic nuclei prior to birth and further suggest that much of the complex structure of the caudal diencephalon is encoded by genetic programs that are active during embryonic development.

**Figure 2.**
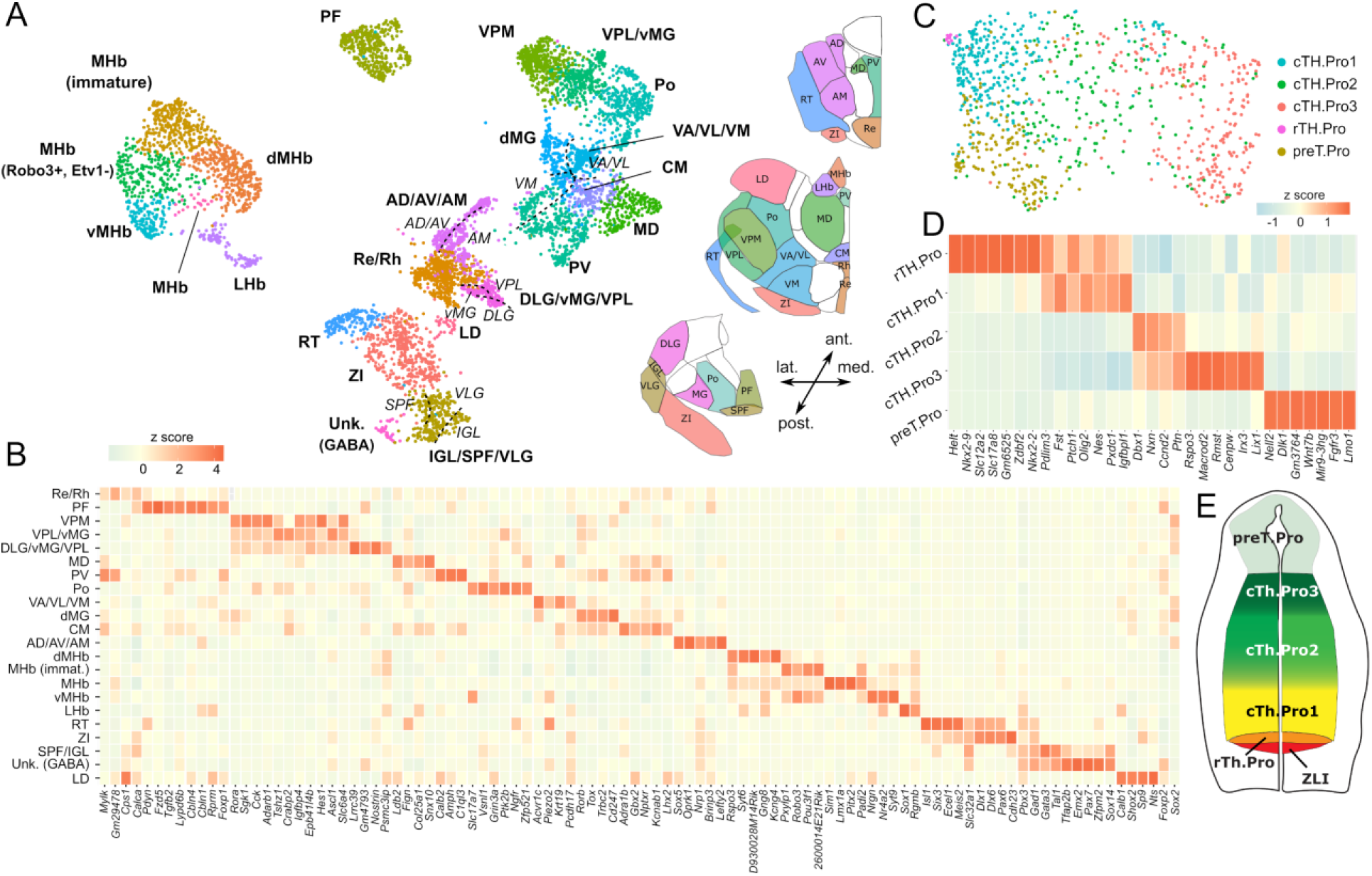
Thalamic nuclei emerge from a diverse pool of progenitor populations and acquire unique transcriptional identities during embryogenesis. **A)** UMAP representation of the mRNA expression data of E18.5 control cells from the post-mitotic cTh.N, rTh.N, ZI, habenula, and reticular complex cell populations. In total, this analysis identified 23 cell subpopulations with distinct transcriptomic profiles (bold labels). Using a spectral graph method, we split some of these populations into smaller transcriptional identities (italic labels). **B)** Heat-map depicting the expression of the top differentially expressed genes in the cell populations from A). **C)** UMAP representation of the mRNA expression data of E12.5 control cells from the *Olig3*^+^ thalamic progenitor population. In total, four thalamic (rTh.Pro and cTh.Pro1-3) and one pretectal (preT.Pro) progenitor populations were identified with distinct transcriptomic profiles. **D)** Heat-map depicting the expression of the top differentially expressed genes in the cell populations identified in C). **E)** Schematic showing the rostro-caudal organization of the identified progenitor cell populations in the developing thalamus as inferred from RNA in situ hybridization data.

### Shared genetic programs across distinct sensory modalities are established during embryogenesis

Sensory inputs in the brain follow cortico-thalamo-cortical loops, where first order thalamic nuclei project peripheral sensory inputs onto the primary sensory cortex, and higher order thalamic nuclei receive their input from the primary sensory cortex and project it back to a secondary cortex ^19,49,50^. The hierarchical position that each nucleus occupies in these circuits has been shown to be the primary determinant of the postnatal transcriptional identity of somatosensory, visual and auditory thalamic nuclei^25^. To assess whether the shared transcriptional programs between same-order nuclei are established earlier during embryogenesis, we projected the postnatal day 3 (P3) differential gene expression signatures of first- and higher-order nuclei from a previous study ^25^ onto our single-cell gene expression data of E18.5 thalamic nuclei. This analysis revealed that postnatal gene expression signatures of first- and higher-order nuclei are already present at E18.5 (Supplementary Fig. 7). We observed high expression of genes that are postnatally associated with first-order nuclei in the VPL, VPM, and vMG cell populations, and of genes that are postnatally associated with higher-order nuclei in the Po, parafascicular (PF), centromedian (CM), medial dorsal (MD), lateral dorsal (LD), and dMG transcriptomic cell subpopulations (Supplementary Fig. 7), in agreement with the hierarchical order of sensory thalamic nuclei in cortico-thalamo-cortical loops ^50^. Of particular note, all of the higher-order sensory nuclei, except for dMG, were negative for *Sox2* and positive for *Foxp2* gene expression (Fig. 2B). This observation suggests that first- and higher-order nuclei emerge to a large extent from different thalamic cell lineages.

### Thalamic progenitors are a heterogeneous cell population

We next characterized the cellular heterogeneity of the *Olig3*^+^ progenitor subpopulation in E12.5 control embryos. Since cell cycle is tightly coupled to cell differentiation during early neurogenesis ^51^, an unsupervised analysis of the progenitor cell population using the most variable genes failed to reveal distinct progenitor types. Regressing out the expression of cell cycle genes did not rectify this issue. To circumvent this problem in the analysis, we adopted a semi-supervised approach in which we compared the transcriptional profile of progenitor cells based on the expression of genes that were highly correlated or anti-correlated with a set of pre-specified markers, including *Nkx2-2*, *Olig2*, *Dbx1*, and *Rspo3*. These genes were previously shown to mark distinct and partially overlapping progenitor domains distributed along the rostral to caudal axis of the thalamus ^30,40^. Clustering and differential expression analysis based on this approach identified five transcriptionally distinct populations of progenitors (Fig. 2C, D, and Supplementary Table 4). We named these progenitor populations as rTh.Pro, cTh.Pro1, cTh.Pro2, cTh.Pro3, and preT.Pro based on their rostro-caudal order in the neural tube (Fig. 2E). The two most rostral thalamic progenitor populations (rTh.Pro and cTh.Pro1) were characterized by the expression of *Shh*-responsive genes, such as *Nkx2-2*, *Olig2*, and *Ptch1* (Fig. 2D), consistent with previous findings ^12^. High levels of expression of Shh responsive genes were excluded from cTh.Pro2 and cTh.Pro3 domains, which were defined by distinct sets of differentially expressed transcripts (Fig. 2D). The most caudal progenitor subpopulation (preT.Pro) was characterized by the expression of pretectal markers (*Pax3, Meis1, Lmo1*) (Supplementary Table 4). On the basis of these results, we hypothesize that different cell lineages derived from distinct thalamic progenitor populations give rise to the diversity of thalamic nuclei.

### Glutamatergic thalamic nuclei emerge sequentially through the coordinated action of three distinct cell lineages

To uncover the transcriptomic lineages that relate thalamic progenitors at E12.5 to post-mitotic thalamic neurons at E18.5, we devised a semi-supervised approach for transferring the E12.5 and E18.5 cell annotations across all time points and identified cells with a similar gene expression profile (Methods). We disaggregated the progenitor and post-mitotic populations into smaller clusters in the full single-cell RNA-seq representation and used the E12.5 and E18.5 cell annotations to label each cluster. We then constructed separate single-cell gene expression representations for glutamatergic cell lineages (*Olig3*^+^ cTh.Pro, cTh.N early post-mitotic, and cTh.N clusters), and computed the RNA velocity vector field ^52,53^ in these representations (Fig. 3A).

**Figure 3.**
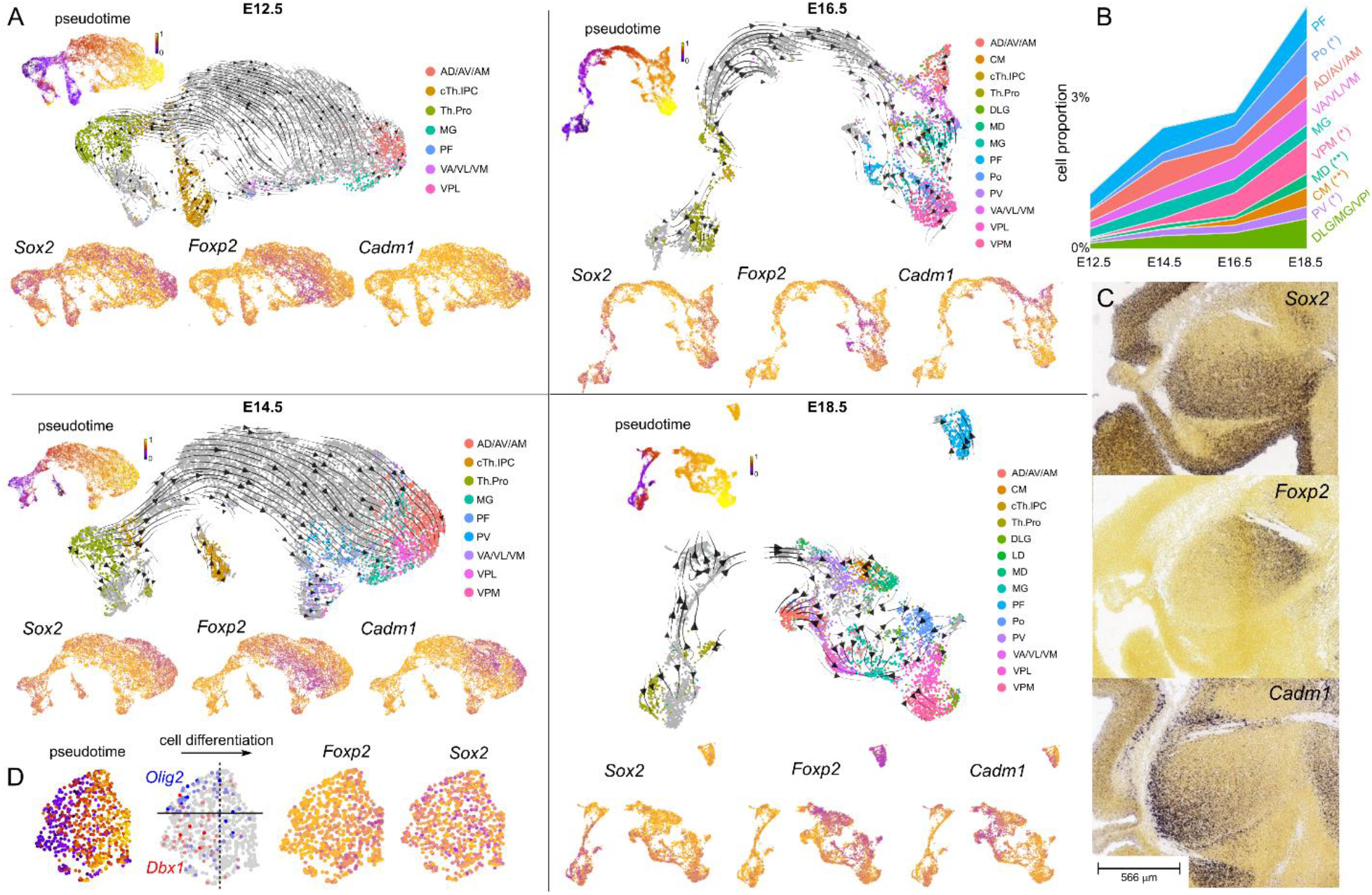
Glutamatergic thalamic cell lineages regionalize the thalamus into distinct molecular domains. **A)** UMAP representation and RNA velocity field of the single-cell RNA-seq data of the glutamatergic cell lineage in control embryos at each developmental stage. For reference, the same UMAP is also colored by the inferred cell differentiation pseudo-time and the gene expression levels of *Sox2*, *Foxp2*, and *Cadm1*. The expression of *Sox2* and *Foxp2* genes separates differentiation trajectories into Sox2^+^ Foxp2^-^ (cTh.N1) and *Sox2*^-^ *Foxp2*^+^ (cTh.N2/3) populations. **B)** Proportion of cells belonging to different glutamatergic thalamic nuclei at each developmental stage. Distinct thalamic subpopulations emerge at different time points. Late- and intermediate- emerging cell subpopulations are indicated by (**) and (*), respectively. **C)** Whole-mount RNA in situ hybridization for *Sox2*, *Foxp2*, and *Cadm1* in sagittal sections of the E13.5 diencephalon. The expression of these markers regionalizes the developing thalamus into distinct domains. Image credit: Allen Institute. **D)** Glutamatergic thalamic IPCs are a heterogeneous population of cells with contributions from both cTh.Pro1 and cTh.Pro2/3 progenitors. The UMAP representation of the single-cell RNA-seq data corresponding to the glutamatergic thalamic IPC population from E14.5 control embryos is colored by the inferred cell differentiation pseudo-time and the gene expression levels of *Olig2*, *Dbx1*, *Foxp2*, and *Sox2*.

Each embryonic day was analyzed independently, since neurodevelopmental lineages may change across time as a consequence of shifts in the progenitor identities. As expected, the inferred differentiation trajectories connected the progenitors with the post-mitotic neurons in each developmental stage (Fig. 3A). The number of *Olig3*^+^ progenitors and intermediate progenitor cells (IPCs) rapidly diminished after E14.5 as a consequence of the transition into gliogenesis, while post-mitotic cells continued to differentiate well beyond E14.5. Based on these data, we conclude that it takes several days for early post-mitotic cells to acquire the unique transcriptional identity of a given thalamic nucleus.

Our analysis revealed the sequential emergence of glutamatergic thalamic nuclei, where lateral and intermediate nuclei such as the AD/AV/AM, DLG/MG/VPL, PF, and Po differentiate first, and medial nuclei such as the CM, MD, and paraventricular (PV) differentiate later (Figs. 3A and B). Differential gene expression analysis along the differentiation trajectories showed that the expression of *Sox2* and *Foxp2* was anti-correlated with each other (Spearman’s correlation coefficient *r* = –0.2, *p*-value < 2 × 10^−16^) and separated glutamatergic cell differentiation trajectories into *Sox2*^+^ *Foxp2*^-^ and *Sox2*^-^ *Foxp2*^+^ trajectories at all developmental stages (Figs. 3A). These results suggest the presence of two major cell differentiation lineages of glutamatergic thalamic neurons, which we denote as cTh.N1 and cTh.N2/3 for reasons described below.

To understand the relationship between these cell differentiation lineages and the observed diversity of thalamic nuclei and progenitors, we examined the expression of *Sox2* and *Foxp2* in these cell populations. The expression of *Sox2* was ubiquitous in the progenitors and was preserved throughout the cTh.N1 lineage, including in the differentiated VPM, AD/AV, AM, VA/VL, and VM nuclei, which originate from this cell lineage (Fig. 3A and Supplementary Fig. 9). By contrast, the expression of *Foxp2* was initiated in a subset of IPCs (Fig. 3D) and preserved throughout the cTh.N2/3 lineage, including the PF, Po, PV, MD, and CM nuclei, which originate from this cell lineage (Fig. 3A and Supplementary Fig. 8). Importantly, these results establish a basis for the distinction between *Sox2*- and *Foxp2*-expressing thalamic nuclei observed in E18.5 cells (Fig. 2B, far right columns).

Our analysis also revealed a separation of the cTh.N1 differentiation trajectories into *Cadm1*^+^ and *Cadm1*^-^ sub-lineages (Fig. 3A). The expression of *Cadm1* was initiated in a subset of early cTh.N1 post-mitotic cells and preserved in the thalamic nuclei that derive from this lineage, including the AD/AV, AM, VA/VL, and VM nuclei (Fig. 3A and Supplementary Fig. 8C). RNA in situ hybridization confirmed the regionalization of the thalamus into three distinct domains based on the expression of *Sox2*, *Foxp2*, and *Cadm1*, with *Sox2* being expressed rostrally and *Foxp2* caudally (Fig. 3C). Thus, our analysis of differentiation trajectories in the single-cell data is consistent with the emergence of glutamatergic thalamic nuclei from distinct transcriptomic lineages marked by the expression of *Sox2* and *Foxp2*. The rostro-caudal organization of these transcriptomic lineages together with the organization of thalamic progenitors in the neural tube suggests that these lineages originate from distinct subpopulations of glutamatergic progenitors.

### Glutamatergic thalamic IPCs are a heterogeneous population of cells with contributions from both cTh.Pro1 and cTh.Pro2/3 progenitors

Despite our ability to transfer annotations across timepoints, a gene expression bottleneck within IPCs prevented us from relating the aforementioned transcriptomic cell lineages to individual thalamic progenitor identities. Most of the genes expressed in progenitor cells ceased to be expressed after mitotic arrest, except for *Hs3st1* which marked both the cTh.Pro1 and cTh.N1 populations (Supplementary Fig. 10). However, a closer look at the IPC population and its RNA velocity field revealed the presence of two subsets of early IPCs, characterized by the expression of the cTh.Pro1 marker *Olig2,* and the cTh.Pro2/3 marker *Dbx1*, as well as two sub-sets of late IPCs, respectively, characterized by the presence and absence of *Foxp2* expression (Fig. 3D). The *Foxp2*^+^ subpopulation of IPCs appeared contiguous in the gene expression space to the *Dbx1*^+^ subpopulation, whereas the *Foxp2*^-^ subpopulation appeared contiguous to the *Olig2*^+^ subpopulation. These results suggest that the cTh.N1 and cTh.N2/3 cell lineages are respectively derived from cTh.Pro1 and cTh.Pro2/3 progenitors, and that both lineages expand through the generation of distinct IPCs.

### GABAergic neurons diverge from progenitors with a shared transcriptional identity

To identify the molecular logic that drives the generation of a diverse pool of inhibitory neurons in the thalamus, we performed an analysis of the GABAergic cell populations similar to the one described above for glutamatergic cell lineages. We constructed separate single-cell gene expression representations for GABAergic cells (GABAergic progenitors, rTh.N, preTh.N early post-mitotic, ZI, and RT clusters) and computed the RNA velocity vector field in these representations for each time point (Supplementary Fig. 9). GABAergic progenitors expressed the bHLH transcription factor *Ascl1* and comprised two major transcriptomic cell lineages that were respectively marked by the expression of *Tal1* and *Helt* (Supplementary Fig. 9). The cell lineage originating from *Helt*^+^ progenitors was characterized by the early expression of *Dlx1* followed by *Pax6* expression. Upon further differentiation, this lineage led to the GABAergic neuronal populations of the ZI and RT (Supplementary Fig. 9). On the other hand, the cell lineage originating from *Tal1*^+^ progenitors split into three distinct sub- lineages that were marked by the expression of *Six3*, *Cbln2*, and *Pax7* (Supplementary Fig. 9). These sub-lineages led respectively to the GABAergic neurons of the IGL/VLG thalamic nuclei, the pretectum, and the tectum. The thalamic sub-lineage was also characterized by the co-expression of Shh-responsive genes, such as *Pdlim3* and *Nkx2-2*, suggesting the involvement of regional Shh signaling in its specification (Supplementary Fig. 9). Although these results are based on the continuity of gene expression programs across cell differentiation and cannot be relied upon to define clonal relationships, they are consistent with recent clonal studies of the mouse forebrain showing a common progenitor origin for GABAergic neurons with drastically different transcriptomic profiles and anatomical locations ^54^. We conclude that GABAergic neurons of the thalamus derive from *Tal1^+^* neural progenitors and are characterized by the expression of Shh-responsive genes.

### Shh signaling is required for rTh.Pro and cTh.Pro1 progenitor specification and expansion

The inference of developmental trajectories from unperturbed single-cell transcriptomic data has proven to be misleading in some situations ^55^. Therefore, we studied the effect of the *ΔSBE1/5* deletions on the transcriptomic cell lineages to further test our model of thalamic development. To determine the effect of Shh signaling on thalamic progenitor specification, we examined the differences between *Olig3*^+^ progenitors in control and *ΔSBE1/5* embryos at E12.5. Our analysis of the single-cell RNA-seq data identified a strong depletion of rTh.Pro and cTh.Pro1 progenitors in *ΔSBE1/5* embryos (Fig. 4A, odds ratio = 13.5, Fisher’s exact test *p*-value < 10^−10^), consistent with previous studies demonstrating that the specification of these two progenitor identities is dependent on Shh signaling ^12,13^. RNA in situ hybridization for *Nkx2-2*, *Tal1*, and *Olig2* confirmed the strong depletion of rTh.Pro and cTh.Pro1 progenitors in *ΔSBE1/5* embryos (Fig. 4B). Our analysis of the single-cell RNA-seq data also identified a moderate expansion of the cTh.Pro2/3 progenitor pool in mutant embryos (Fig. 4A, odds ratio = 0.23, Fisher’s exact test *p*-value < 10^−10^). However, we were unable to confirm this expansion with in situ data (Fig. 4B). Differential gene expression analysis between control and *ΔSBE1/5 Olig3*^+^ glutamatergic progenitor cells revealed the downregulation of cell cycle genes in mutant cells (Fig. 4C, Supplementary Fig. 11, and Supplementary Table 1). This effect was particularly prominent in the IPC population (Fig. 4C). We confirmed the observed downregulation of cell cycle in *ΔSBE1/5* cTh.Pro1 progenitors by means of a 5-ethynyl-2’-deoxyuridine (EdU) incorporation assay (Fig. 4D, E, and Supplementary Fig. 12). These results suggest that the specification and expansion of cTh.Pro1 progenitors is dependent on Shh. Moreover, consistent with our hypothesis that cTh.N1 cells derive from cTh.Pro1 progenitors, we observed that the reduction in cTh.Pro1 progenitors in *ΔSBE1/5* mice was accompanied by a substantial reduction in the number of *Sox2*^+^ post-mitotic cells in the glutamatergic cTh.N1 transcriptomic lineage in our single-cell RNA-seq data and a higher proportion of *Foxp2* expressing cells in this lineage (Fig. 4F and Supplementary Fig. 13). These results were confirmed by immunofluorescence staining for Sox2 and Foxp2 in control and *ΔSBE1/5* embryos at E14.5 (Fig. 4G, H). We conclude that Shh signaling is required for the specification and expansion of rTh.Pro and cTh.Pro1- derived thalamic cell lineages.

**Figure 4.**
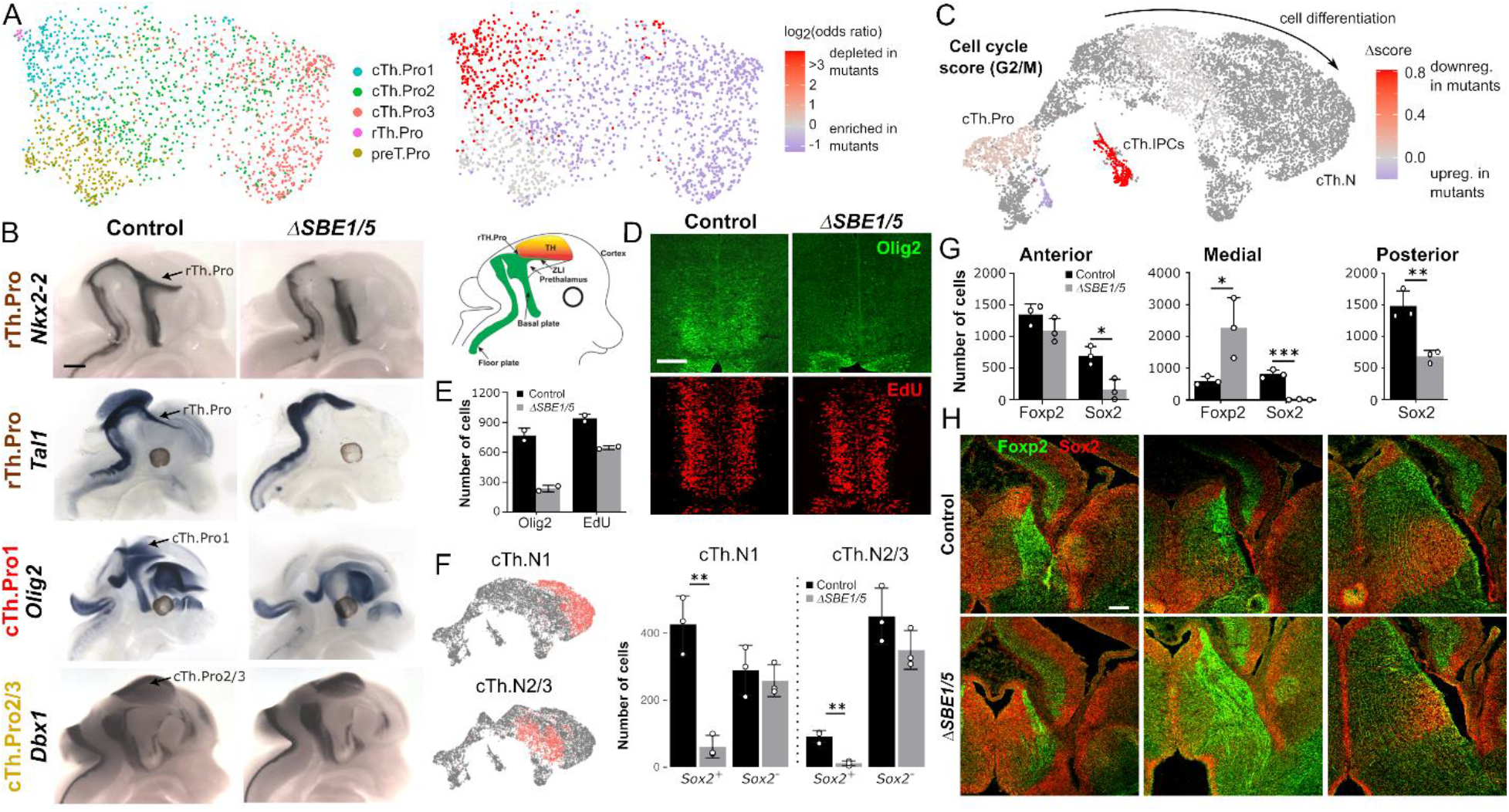
Shh signaling is required for the specification and expansion of rTh.Pro and cTh.Pro1 thalamic progenitors. **A)** UMAP representation of the mRNA expression data of E12.5 control and *ΔSBE1/5* cells from the *Olig3*^+^ thalamic progenitor population (left). The UMAP is colored by the depletion or enrichment of cells between control and *ΔSBE1/5* embryos for each progenitor subpopulation, showing a strong depletion of rTh.Pro and cTh.Pro1 cells in *ΔSBE1/5* embryos. **B)** Whole-mount RNA in situ hybridization for *Nkx2-2*, *Tal1*, *Olig2*, and *Dbx1* in E12.5 control and *ΔSBE1/5* embryos, confirming the strong depletion of rTh.Pro and cTh.Pro1 cells in mutant mice (n=3) Scale bar = 500 μm. **C)** UMAP representation of the glutamatergic thalamic cell lineage at E14.5 colored by the difference in the G2/M cell cycle gene expression score between control and *ΔSBE1/5* cells. The analysis shows G2/M cell cycle genes are downregulated in thalamic progenitors and IPCs in *ΔSBE1/5* embryos. **D, E)** EdU incorporation and immunofluorescence staining for Olig2 on coronal sections of the thalamus in control and *ΔSBE1/5* E13.5 embryos, showing a depletion of Olig2 expressing cells (cTh.Pro1 cells) and a reduction of cell cycle in IPCs (n=2). Scale bar = 100 μm. **F)** The total number of *Sox2* expressing cells in the post-mitotic cTh.N1 and cTh.N2/3 transcriptomic cell lineages is largely depleted in *ΔSBE1/5* compared to control embryos at E14.5. For reference, the location of the post-mitotic cTh.N1 and cTh.N2/3 clusters in the UMAP representation of the glutamatergic thalamic cell lineage is shown on the left. **G, H)** Immunofluorescence staining for Sox2 and Foxp2 in anterior, medial, and posterior sections of E14.5 control and *ΔSBE1/5* embryos. A depletion of Sox2^+^ cells and an expansion of Foxp2^+^ cells is observed in mutant embryos (*p<0.05, **p<0.01, ***p<0.001, Student’s t-test, n=3, error bars represent standard deviation). Scale bar = 100 μm.

### *Nkx2-2* and *Sox2* expressing thalamic nuclei are reduced in *ΔSBE1/5* mice

Based on our model of thalamic development, the observed depletion of cTh.Pro1 progenitors in *ΔSBE1/5* embryos should lead to a failure in the development of *Sox2*-expressing cTh.N1 thalamic nuclei. Our analysis of single-cell RNA-seq data from post-mitotic glutamatergic cells in control and *ΔSBE1/5* embryos at E18.5 confirmed a large depletion of cells in thalamic nuclei expressing high levels of *Sox2*, including the AD/AV/AM, VA/VL/VM, DLG/MG/VPL, and VPM nuclei (Fig. 5A). Consistent with these results, RNA in situ hybridization for some of the markers identified in our differential gene expression analysis of these nuclei (Fig. 2B) showed a large reduction in the size of these nuclei in newborn (P0) *ΔSBE1/5* mice (Fig. 5B and Supplementary Fig. 14). In particular, the AD/AV/AM and VM nuclei were completely absent in mutant mice, whereas the VA/VL, DLG/MG/VPL, and VPM were largely reduced (Fig. 5B and Supplementary Fig. 14). Additionally, Nkx2-2 immunostaining revealed the absence of the IGL/VLG nuclei in *ΔSBE1/5* mice at P0 (Supplementary Fig. 14), consistent with the depletion of GABAergic rTh.Pro progenitors (Fig. 4A). Taken together, the analysis of single-cell RNA-seq and in situ expression data is consistent with a model of thalamic development where glutamatergic *Sox2*-expressing nuclei (cTh.N1 lineage) are derived from Shh-dependent cTh.Pro1 progenitors, glutamatergic *Foxp2*-expressing nuclei (cTh.N2/3 lineage) are derived from Shh-independent cTh.Pro2/3 progenitors, and GABAergic *Nkx2-2*-expressing nuclei (rTh.N lineage) are derived from *Shh*-dependent rTh.Pro progenitors.

**Figure 5.**
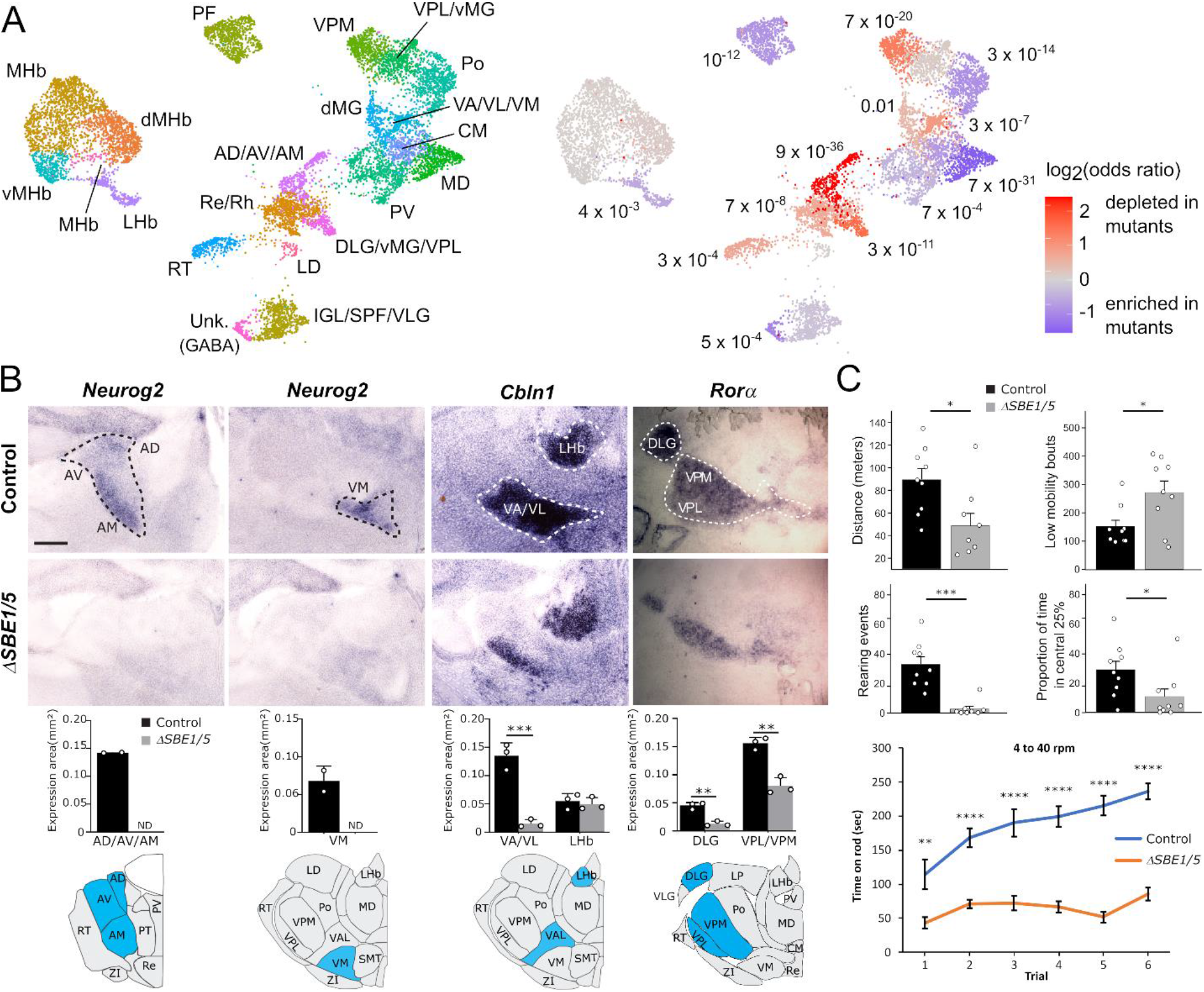
Shh-dependent thalamic nuclei fail to develop in *ΔSBE1/5* mice. **A**) UMAP representation of the single-cell RNA-seq expression data from control and *ΔSBE1/5* embryos at E18.5 corresponding to post-mitotic cTh.N, rTh.N, ZI, habenula, and reticular complex cell populations (left). The UMAP is colored by the amount of depletion or enrichment in the number of cells of each progenitor subpopulation between control and *ΔSBE1/5* embryos (right). **B)** Whole-mount RNA in situ hybridization for *Neurog2*, *Cbln1*, and *Rorα* in coronal sections of the thalamus from control and *ΔSBE1/5* mice at P0, showing a large reduction in the size of the AV/AM/AD, VA/VL/VM, DLG and VPM/VPL thalamic nuclei, consistent with the results of the single-cell RNA-seq analysis. **C)** Locomotor behavior is compromised in *ΔSBE1/5* mice as determined by force plate actomoter (top four graphs) and rotarod (bottom) assays (*p<0.04, **p<0.004, ***p<0.0001, ****p<0.00001, two-tailed t test; n = 9 for force plate; n=10 for rotarod; error bars represent SEM).

### Locomotor deficits in *ΔSBE1/5* mice

We next assessed the consequences that abnormal thalamic development might have on animal behavior in *ΔSBE1/5* mutant mice. *ΔSBE1/5* mice are viable and exhibit reduced locomotor activity and rearing in the open field, as well as a significantly impaired initial coordination and subsequent motor learning on the accelerating rotarod test (Fig. 5C). These early onset motor deficits resemble several cardinal features of infantile Parkinson’s disease, including bradykinesia, abnormal gait, and poor coordination (Fig. 5C) ^56^.

The nigrostriatal pathway is a critical component of the basal ganglia motor circuit that degenerates in individuals with Parkinson’s disease. However, unlike other mouse models of the infantile form of this condition that result from tyrosine hydroxylase deficiency ^56^, the rate limiting enzyme in dopamine synthesis, we observed no defects in the development or maintenance of dopaminergic neurons in the substantia nigra pars compacta or their projections to the striatum in *ΔSBE1/5* mice (Supplementary Fig. 15). This result is particularly relevant since SBE1 and SBE5 also regulate *Shh* expression in the ventral midbrain (Fig. 1A), which is the source of midbrain dopaminergic neurons. However, since these neurons develop properly in *ΔSBE1/5* mice, likely due to their dependency on an earlier source of Shh, we suspect that another component of the basal ganglia motor circuit is compromised in *ΔSBE1/5* mice.

For several reasons, we favor the loss of motor thalamic nuclei (VA/VL, VM) as a likely explanation for the locomotor deficits in *ΔSBE1/5* mice. Firstly, as described above, the spatiotemporal regulation of *Shh* expression in the ZLI is critical for the elaboration of cTh.Pro1 that populates multiple thalamic nuclei, including VA/VL and VM. Secondly, depletion of cTh.Pro1 results in a profound reduction of motor thalamic nuclei in *ΔSBE1/5* mice. Thirdly, lesions to VA/VL cause severe motor dysfunction in monkeys, rats and mice, further highlighting the importance of the motor thalamus in relaying excitatory signals to the motor cortex ^57–60^. Taken together, these results illustrate the utility of deciphering pathogenic mechanisms of thalamic dysfunction in *ΔSBE1/5* mice as a means to gain novel insight into the etiology of neurodevelopmental disorders, such as infantile Parkinson’s disease.

## Discussion

Our study provides a comprehensive transcriptome-wide analysis of thalamic progenitors and their trajectories into thalamic nuclei during embryonic stages of brain development. We demonstrate that molecular signatures of thalamic neuronal subtypes can be readily distinguished as early as E12.5, prior to their aggregation into histologically distinct thalamic nuclei ^28^. These data lend support to the “outside-in” model of thalamic neurogenesis, whereby early born neurons contribute to lateral thalamic nuclei and later born neurons contribute to medial thalamic nuclei ^34,61^. Our work further extends this model with molecular insights into the mechanisms by which diverse thalamic nuclei acquire their identities. By following single-cell trajectories over developmental time, we show that Shh responsive cTh.Pro1 progenitors give rise to glutamatergic neurons (cTh.N1) within sensory (DLG, VPM/VPL, vMG), motor (VA/VL, VM) and anterior (AD, AV, AM) nuclei that occupy predominantly ventrolateral regions of the thalamus. Shh responsive rTh.Pro progenitors were shown to give rise to GABAergic neurons (rTh.N) in VLG and IGL thalamic nuclei, in agreement with previous findings ^12,13^. In contrast, Shh independent cTh.Pro2/3 progenitors give rise to neurons (cTh.N2/3) within nuclei (e.g. PV, MD, CM) that form at more dorsomedial positions of the thalamus. Not all medial thalamic nuclei (e.g. CL, LP, PCN, PT, SMT) were readily discerned in our scRNA-seq analysis, possibly due to limitations in our ability to detect their unique transcriptional signatures, or because their maturation extends beyond the prenatal period of brain development.

The bifurcation of lineage commitments according to thalamic progenitor identity is in general agreement with results from in vivo clonal analyses ^33,34^. These studies revealed that individual thalamic progenitors give rise to many neurons that populate multiple thalamic nuclei. They also demonstrated that the clonal relationship between thalamic nuclei is determined primarily by the rostro-caudal and dorso-ventral positions of thalamic progenitors. These findings are consistent with our observations that thalamic nuclei originate from a small number of spatially and temporally segregated neural progenitors located at key positions along the primary axes of the developing thalamus (Fig. 6).

**Figure 6.**
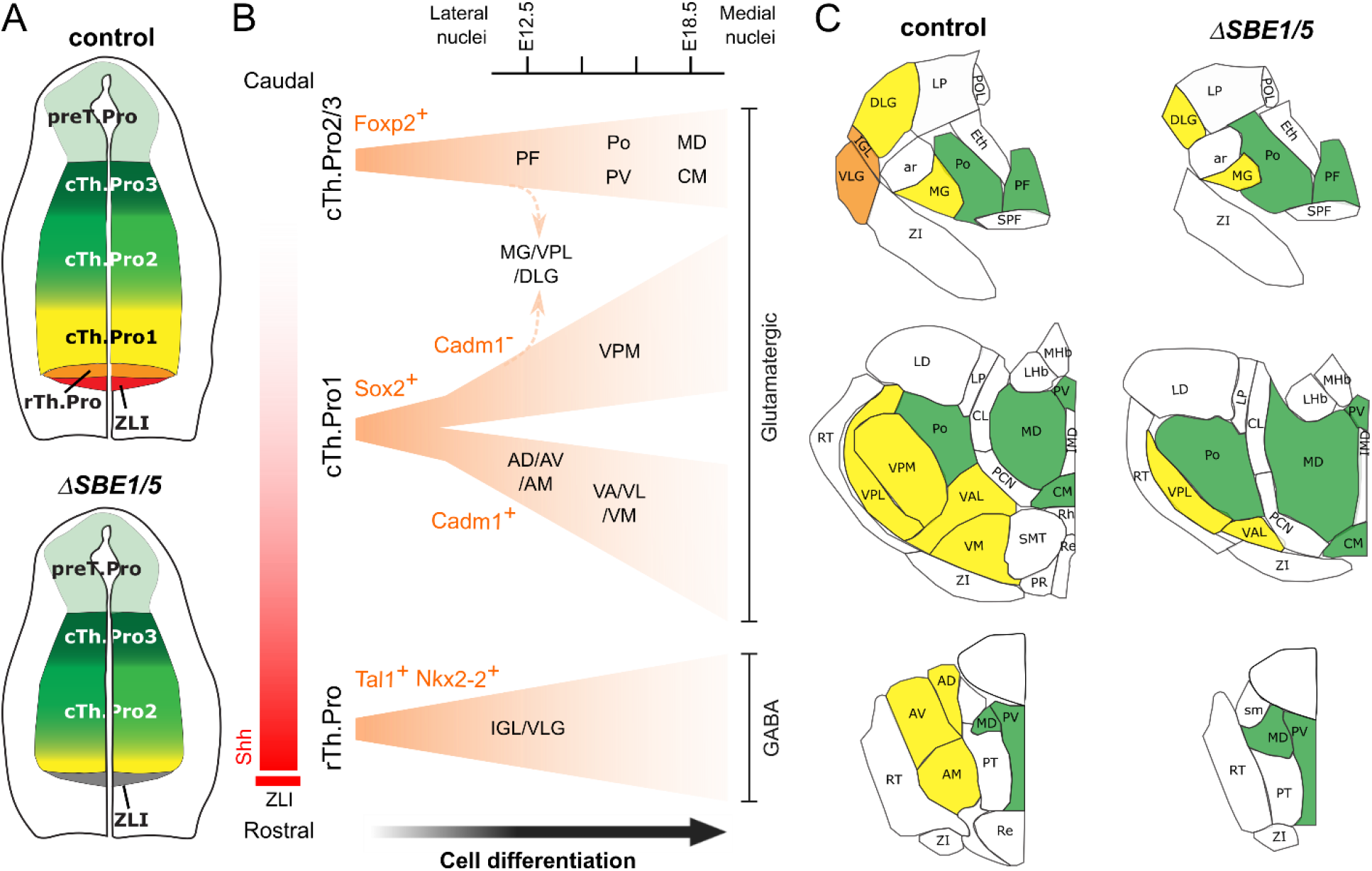
Model of thalamic development. **A)** Thalamic progenitors are organized rostro-caudally in the neural tube and acquire distinct identities based on their exposure to the Shh morphogen gradient originating from the ZLI and basal plate (top). In *ΔSBE1/5* mutants, the pool of rTh.Pro and cTh.Pro1 progenitors are not specified and fail to expand due to the lack of Shh signaling (bottom). **B)** rTh.Pro, cTh.Pro1, and cTh.Pro2/3 give rise to different cell lineages which are characterized, respectively, by the expression of *Tal1*, *Sox2*, and *Foxp2* at all post-mitotic stages of cell differentiation. The *Sox2*^+^ lineage is divided into two sub-lineages soon after post-mitotic arrest of the progenitors. These sub-lineages can be distinguished by the expression of *Cadm1*. Different cell lineages contribute to different thalamic nuclei in a sequential temporal manner, where lateral nuclei are formed first and medial nuclei are formed later. An exception to this model is the MG/VPL/DLG, which receives contributions from both cTh.Pro1 and cTh.Pro2/3 progenitors. **C)** Thalamic nuclei derived from Shh-dependent (rTh.Pro and cTh.Pro1) progenitors (labelled in orange and yellow, respectively) fail to develop in *ΔSBE1/5* mice, whereas nuclei derived from cTh.Pro2/3 progenitors (labelled in green) are partially expanded in the absence of Shh.

Consistent with this model, our scRNA-seq analysis also identified heterogeneity in the IPC lineage. Previous work described the presence of IPCs in the thalamus but not their specific lineage relationships with thalamic progenitors and neurons ^40,62^. Our data reveal that Olig2^+^ Sox2^+^ IPCs give rise to ventrolateral thalamic nuclei and Foxp2^+^ IPCs give rise to dorsomedial thalamic nuclei (Fig. 6). Intermixing of Sox2^+^ and Foxp2^+^ lineages was observed in a subset of sensory nuclei (Fig. 6B). The Sox2^+^ lineage is further partitioned by *Cadm1* expression. In addition to marking distinct and overlapping thalamic IPCs and neurons, Sox2 and Foxp2 are required to regulate functional properties of their respective thalamic lineages ^63,64^. In particular, the cell autonomous loss of VP (VPM/VPL), PF and Po nuclei in *Foxp2* mutants at E14.5 ^63^, suggests that Foxp2 may be required for the transition of IPCs to a subset of cTh.N2/3 thalamic neurons. It is worth noting that Foxp2 has a similar role in the cerebral cortex where it regulates the formation of IPCs and their transition to cortical neurons ^65^. Less is known about the role of Sox2 in thalamic neurogenesis. However, the conditional knockout of Sox2 in post-mitotic thalamic neurons results in a reduction in the size and connectivity of sensory nuclei ^64^. Thus, the developmental segregation of cTh.Pro, IPC and cTh.N subtypes by lineage-determining transcription factors likely plays an important role in the formation of distinct thalamic nuclei.

The transcriptional profiles of thalamic nuclei at early postnatal stages were previously shown to be more discriminative when classified based on hierarchical order rather than by sensory modality ^25^. For instance, first order (VPM, DLG, vMG) and higher order (Po, LP, dMG) thalamic nuclei display a greater degree of gene expression overlap within their respective hierarchical order than do pairs of nuclei that function in similar somatosensory (VPM/Po), visual (DLG/LP) and auditory (vMG/dMG) pathways. Moreover, the gene expression differences between first order and higher order nuclei become more pronounced as first order nuclei increase their peripheral connections to primary sensory cortical targets at early postnatal stages (P3-P10). However, when we examined first order and higher order gene expression signatures in our scRNA-seq dataset, we were surprised to find that several thalamic nuclei already displayed transcriptional profiles characteristic of their hierarchical order at E18.5 (Supplementary Fig. 7), a stage when thalamocortical synapses are still relatively immature. Therefore, to reconcile these differences, we suggest that the input-dependent logic of first order and higher order thalamic nuclei acts on a pre-specified transcriptional program that initiates at embryonic stages of thalamic development and continues to be refined into the early postnatal period. As neuronal connections between thalamic nuclei and their cortical and subcortical targets continue to mature, additional gene expression networks are likely to emerge that further distinguish differences or consolidate similarities between thalamic nuclei ^41,66^.

Deciphering the pathogenic mechanisms of thalamic dysfunction in *ΔSBE1/5* mutants provided novel insight into the etiology of a neurodevelopmental movement disorder with a similar phenotype to infantile Parkinson’s disease. We demonstrate that the spatiotemporal regulation of *Shh* expression in the ZLI and basal plate of the caudal diencephalon is critical for the elaboration of thalamic progenitor identities that populate multiple thalamic nuclei, including the motor thalamus (VA/VL, VM), principal sensory nuclei (DLG, VPM/VPL, vMG) and anterior thalamic nuclei (AD, AV, AM) (Fig. 5). It would therefore appear that the timing of Shh depletion may explain many of the unique features of our mouse model compared to other conditional *Shh* mutants ^11,12^.

The basal ganglia are subcortical nuclei that control volitional movement, as well as non-motor functions, and have been implicated in Parkinson’s disease and other movement disorders ^67–70^. The prevailing model of basal ganglia function stipulates that distinct populations of medium spiny neurons in the striatum facilitate and suppress motor behavior via direct and indirect pathways, respectively ^71^. Specifically, direct pathway medium spiny neurons (dMSN) promote movement by reducing the inhibitory effects of major output nuclei, internal globus pallidus and substantia nigra pars reticulata, on the motor thalamus (VA/VL, VM), resulting in the relay of excitatory motor signals to the primary motor cortex. In contrast, medium spiny neurons of the indirect pathway (iMSN) suppress thalamocortical circuitry and movement by strengthening repressive inputs from the external globus pallidus and subthalamic nucleus on the motor thalamus.

Motor impairment in Parkinson’s disease is caused by the degeneration of dopamine expressing neurons in the substantia nigra pars compacta, favoring activation of iMSNs over dMSNs ^71^. Shh has been implicated in the production and survival of dopaminergic neurons in the nigrostriatal pathway ^72–75^. However, since midbrain dopaminergic neurons are unaffected in *ΔSBE1/5* mutants, other pathogenic mechanisms must be invoked to explain the motor deficits in these mice.

Recent studies have also implicated the thalamus in Parkinson’s disease pathology ^76–78^. In one report, dopamine depletion altered synaptic strength of thalamo- striatal circuits, shifting the balance from direct to indirect pathways and thereby reducing movement ^77^. In another study, optogenetic photostimulation of inhibitory basal ganglia inputs from the globus pallidus caused post inhibitory rebound firing of ventrolateral (VL) thalamic neurons that induced muscle contractions, rigidity, tremors, and hypo-locomotor activity ^79^. Lesions to VA/VL cause severe motor dysfunction in monkeys, rats and mice, further highlighting the importance of motor thalamic nuclei in relaying excitatory signals to the motor cortex ^57–60^.

We propose that the motor impairment in *ΔSBE1/5* mutants is attributed to loss of the Shh-dependent cTh.Pro1 subtype of thalamic progenitors, resulting in a reduced number of Olig2^+^ Sox2^+^ IPCs and subsequently, fewer cTh.N1 thalamic neurons populating motor thalamic nuclei (VA/VL, VM) during embryonic development. Future studies will address the consequences that alterations of other thalamic nuclei have on sensory, motor and other behaviors in *ΔSBE1/5* mutant mice. These experiments have the potential to further improve our basic understanding of thalamic development, which has consistently lagged behind other brain regions, and may also provide novel insights into the etiology of other circuit-level endophenotypes associated with abnormal motor and sensory information processing that occur in a variety of neurodevelopmental disorders ^80–83^.

## Acknowledgments

The authors are grateful to Drs. Stephen Liebhaber, Zhaolan Zhou, and Hao Wu for their constructive comments on the manuscript. They also thank the Center for Applied Genomics of the Children’s Hospital of Philadelphia for excellent technical assistance with the preparation and sequencing of single-cell RNA-seq libraries. This work was supported by NIH grant R01 NS039421 (DJE).

## Author contributions

K.G. performed the computational analyses of the single-cell RNA-seq data. S.W. assisted with the preliminary computational analyses of the single-cell RNA-seq data. S. C. and P.S. performed thalamic dissections and prepared single cell suspensions of thalamic tissue. S.C, T.C. and Y.Y. performed gene expression studies on thalamic tissue. P.S. performed behavioral studies in conjunction with M.F. D.J.E. and P.G.C. jointly supervised the work and wrote the manuscript.

## Methods

### Mouse lines

All mouse experiments were performed in accordance with the ethical guidelines of the National Institutes of Health and with the approval of the Institutional Animal Care and Use Committee of the University of Pennsylvania. Mice were housed in Thoren caging units under a constant 12-hour light/dark cycle. Mice carrying targeted deletions of SBE1 and SBE5 were described previously ^13,42^. To generate *Shh^ΔSBE1ΔSBE5/ΔSBE1ΔSBE5^* double homozygous mutant embryos, the *Shh^ΔSBE5/+^* line was first crossed with *Shh^ΔSBE1/ΔSBE1^* mutants. *Shh^ΔSBE1/+; ΔSBE5/+^* males, carrying the SBE1 and SBE5 deletions in *trans*, were then bred to wild type CD1 females. The progeny from this cross were screened for recombination events that placed the SBE1 and SBE5 deletions in *cis* (1/600 offspring). The *Shh^ΔSBE1ΔSBE5/+^* double heterozygous animals were then intercrossed to generate *Shh^ΔSBE1ΔSBE5/ΔSBE1ΔSBE5^* double homozygous mice and embryos. The 429M20eGFP BAC transgenic mouse reporter line (referred to herein as *Shh-GFP*) expresses eGFP in the ZLI and basal plate of the caudal diencephalon under the transcriptional control of SBE1, as described previously ^84^.

### In situ hybridization

Embryonic or neonatal (P0) brains were collected from timed pregnant females (vaginal plug = E0.5). For whole-mount RNA in situ hybridization, heads were fixed in 4% paraformaldehyde at 4°C for overnight, bisected along the mid-sagittal plane and hybridized with digoxygenin-UTP-labeled riboprobes as previously described ^84^. For RNA in situ hybridization on sections, heads were dissected and fixed for 2 hours in 4% paraformaldehyde at 4°C, then washed in PBS. Samples were cryoprotected overnight in 30% sucrose/PBS then snap frozen in OCT embedding compound (Sakura Finetek Torrence, CA). Samples were serially sectioned along the coronal plane at 16 µm (for E12.5 and E13.5 embryos), 18 µm (for E14.5 embryos) or 20 µm (for E18.5 embryos) thickness using a cryostat (Leica Biosystems, CM3050 S). Sections were hybridized with digoxigenin- UTP-labeled riboprobes as previously described ^85^.

### Immunohistochemistry

Brains were processed for immunohistochemistry in the same fashion as for in situ hybridization on sections. Brain sections were stained with DAPI and incubated with the following primary antibodies: rabbit anti-Foxp2 (1:200, abcam, ab16046), mouse anti- Nkx2.2 (1:300, DSHB, 74.5A5), rabbit anti-Olig2 (1:300, Millipore, AB9610), mouse anti- Sox2 (1:100, R&D Systems, MAB2018), rabbit anti-Sox2 (1:300, Chemicon, Cat#AB5603), and rabbit anti-Tyrosine Hydroxylase (1:1000, Pel-Freez, P40101-0). Detection of primary antibodies was achieved using secondary antibodies conjugated to Goat anti-mouse Alexa488 (1:400, Thermo Fisher, Cat#A28175), Goat anti-rabbit Alexa488(1:400, Thermo Fisher, Cat#A-11008), and Goat anti-rabbit Alexa594 (1:400, Thermo Fisher, Cat#A-11037). Specimens were imaged on a Leica TCS SP8 MP system.

### EdU incorporation

EdU was dissolved in sterile water and administered to pregnant dams via intraperitoneal injection at a concentration of 50 µg/g of body weight, 2 hours prior to embryo harvest (Molecular Probes). Embryos were fixed in 4% paraformaldehyde at 4°C for 2 hours, then were processed in the same fashion as for in situ hybridization on sections. EdU incorporation was detected at room temperature with the Click-iT® EdU Imaging Kit (Molecular Probes #C10339). The staining protocol was optimized for frozen sections using the following modifications: 2x 10’ PBS-Tween wash, 2x 10’ 3% BSA incubation, 30’ incubation in the dark with Click-iT® reaction cocktail assembled in the recommended order immediately prior to application, 3% BSA wash, 2x PBS wash.

### Quantification and statistical analysis of imaging data

All cell counts were performed using the cell counter function in ImageJ (NIH) on tissue sections from at least three control and mutant embryos. In cases where double labeling was examined the tissue was imaged at a single Z-plane. Each channel (green for marker 1, red for marker 2) was first examined independently, assigning a positive count for a given marker to the DAPI stained nucleus most closely associated with the staining. A cell was only counted as double labeled if a single nucleus marked by DAPI had been assigned to the cell labeled by marker1 and marker 2. Statistical analysis of all cell counts was performed in GraphPad Prism using the Student’s *t*-test. For a given in situ probe, expression area was measured from at least three control and mutant embryos using ImageJ software. Quantification of the spatial distribution of genes expressed in the zli was normalized to head size. Statistical analysis of all area and length measurements was performed in GraphPad Prism using the Student’s *t*-test.

### Isolation of embryonic thalamus and single-cell dissociation

The thalamus was manually dissected from control (*Shh^ΔSBE1;ΔSBE5/+^*) and mutant (*Shh^ΔSBE1;ΔSBE5/ΔSBE1;ΔSBE5^*) brains at four embryonic stages (E12.5, E14.5, E16.5 and E18.5) in ice-cold PBS. The caudal and rostral boundaries of dissection coincided with the cephalic flexure and the mammillary body, respectively. Each thalamus was cut into small pieces and dissociated into a single cell suspension using the Papain Dissociation System (Worthington, LK003153) according to the manufacturer’s protocol. Samples were dissociated in Papain-EBSS solution for 40 minutes at 37°C. Papain was inactivated with ovomucoid protease inhibitor, and the digested tissue was resuspended in PBS (calcium and magnesium free) containing 0.04% weight/volume BSA (400 µg/ml). Cell suspensions were stained with Trypan Blue to determine the ratio of viable to damaged cells and counted using a haemocytometer (Thermo Fisher Scientific). Single cell suspensions containing more than 90% viable cells were fixed in methanol for 1 week. Samples were rehydrated at a concentration of 700-1200 cells/ml in ice-cold PBS containing 0.04% weight/volume BSA (400 µg/ml) immediately prior to the generation of single-cell RNA sequencing libraries.

### Single-cell RNA library preparation and sequencing

Single-cell RNA-seq libraries were generated using the 10X Genomics platform (Chromium Single Cell 3’ library and Gel Bead Kit v3, PN-1000075) and sequenced on a NovaSeq 6000 system (Illumina) at the Center for Applied Genomics (Children’s Hospital of Philadelphia). Three independent libraries were generated for each time point and genotype (*n*=24 libraries).

### Processing of RNA sequencing data

Fastq files were aligned to the mm10 mouse reference genome and count matrices were generated using the CellRanger (v2.1) pipeline. Except where otherwise specified, we processed and visualized the scRNA-seq counts with the following Seurat-based pipeline, using Seurat v3.0.2 ^86^. We first scaled and centered the UMI counts. We used the default vst method to identify the top 2,000 variable genes, removing all genes from the X and Y chromosomes to reduce the effect of unequal male and female mouse replicates between conditions. To correct for non-biological batch effects between conditions and time points, we used the Harmony algorithm ^87^ with its Seurat integration, run on the top principal components (PCs) of the variable genes. Harmony outputs a batch-corrected representation of the scRNA-seq data of same dimensionality as the input PCs. We ran Louvain clustering and UMAP on the output of Harmony to visualize this consolidated gene expression space. The full dataset was visualized using 20 PCs and 3 UMAP dimensions, while the rest of the subset analyses used 20 PCs and 2 UMAP dimensions. To compute differentially expressed genes (DEGs) in the clusters, we used edgeR’s generalized linear model likelihood ratio test (glmLRT) ^88^ to compare the gene expression in cells from control mice in a single cluster versus all other clusters in the representation. We also computed DEGs between cells from control and *ΔSBE1/5* mice in each cluster using the same method. To reduce the running time of edgeR, we subsampled large clusters to 2,000 cells when computing DEGs.

### Annotation of cell populations in the full atlas

We followed the steps outlined above (*Processing of RNA sequencing data*) to visualize and cluster all cells in our scRNA-seq dataset. We used the DEGs in each cluster to annotate it based on cell type, differentiation stage, or area of the brain according to published literature and ISH images from the Allen Developing Mouse Brain Atlas.

### Identification of thalamic nuclei at E18.5

We selected all E18.5 cells from the clusters that we annotated as cTh.N, rTh.N, RT, ZI, and habenula and followed the steps outlined above (*Processing of RNA sequencing data*) to visualize and cluster differentiated cells from thalamic nuclei. We compared DEGs from each cluster to known markers of thalamic nuclei and E18.5 mice ISH data from the Allen Developing Mouse Brain Atlas. We used RayleighSelection ^48^ to identify significantly localized genes marking nuclei that could not be disaggregated by unsupervised clustering. Enrichment of each annotated thalamic nucleus in control or *ΔSBE1/5* mice compared to the rest of the E18.5 thalamic clusters was computed using a Fisher’s exact test.

### Identification of progenitor populations at E12.5

We selected all E12.5 cells from the progenitor cell clusters with *Olig3* expression: cTh.Pro, GABAergic progenitors, and astroglia. We used the same approach described above (*Processing of RNA sequencing data*) to produce higher resolution clusters of just these cells and selected those clusters that had high expression of *Olig3* and *Vim* but did not yet express neuronal differentiation markers (*Neurod1*, *Stmn2*). We created a UMAP visualization of the resulting 1,885 cells using the top 10 genes that were correlated and the top 10 genes that were anti-correlated with *Nkx2-2*, *Olig2*, *Dbx1*, and *Rspo3* (62 genes in total). We associated each progenitor subpopulation (rTh.Pro, cTh.Pro1, cTh.Pro2, cTh.Pro3, cTh.IPC, or preT.Pro) with a subset of these 62 genes based on known markers and a correlation-based hierarchical clustering of the genes. We then assigned each cell to a progenitor subpopulation based on the total counts for each set of genes. Enrichment of each subpopulation in control or *ΔSBE1/5* mice compared to the rest of the E12.5 progenitor clusters was computed using a Fisher’s exact test.

### Transferring annotations across time points

We transferred thalamic nuclei and progenitor identities across time points by building representations from the clusters cTh.N, rTh.N, RT, ZI, and habenula (for thalamic nuclei), and cTh.Pro, GABAergic progenitors, and astroglia (for progenitors). We processed and clustered these representations following the same approach as in *Processing of RNA sequencing data*. We annotated each cluster, containing cells from all time points, based on the proportion of annotated E18.5 or E12.5 cells, as long as they represented at least 2% of the cells in the cluster. To identify Shh-responsive clusters, we calculated the GSEA score of Shh-responsive genes (*Gli1*, *Ptch1*, *Olig2*, *Nkx2-2*, *Pdlim3*, *Fst*, *Zdbf2*, *Hs3st1*, and *Slc38a11*) in the list of all genes ordered by fold change in expression between control and *ΔSBE1/5* mice.

### Thalamic lineages across time

We reconstructed the GABAergic and glutamatergic thalamic lineages separately for each time point. For GABAergic lineages, we selected cells from the GABAergic progenitors, rTh.N (early post-mitotic), rTh.N, preTh.N (early post-mitotic), RT, and ZI clusters, as well as the cells labelled as rTh.Pro at E12.5 (*Identification of progenitor populations at E12.5*). For glutamatergic lineages, we selected cells from the cTh.N and cTh.N (early post-mitotic) clusters, as well as progenitors from all time points in *Olig3^+^* progenitor clusters. We prepared UMAP representations of these differentiation lineages as described in *Processing of RNA Sequencing data*, with the addition of Seurat’s cell cycle regression to balance out the clear cell cycle effect differences between progenitor and differentiated cells. Cell cycle scores were computed using Seurat’s CellCycleScoring function. We used the velocyto command line interface ^52^ and scVelo ^53^ to infer and visualize RNA velocity streamlines and pseudotime on the UMAP representations. We identified gene patterns associated with the differentiation trajectory from the top DEG lists, then further identified correlated and anti-correlated genes of interest. The significance of a change in Seurat’s G2/M score between control and *ΔSBE1/5* cells in each cluster was calculated using a Wilcoxon rank-sum test.

### Locomotor Behavior

Locomotor activity was assayed using the force plate actometer as previously described ^89^. Briefly, mice (10 to 14 weeks of age) were acclimated to the room for 15 minutes prior to the start of each experiment. Individual mice were placed on an open field plate (28 cm × 28 cm) with four force transducers and sampled at 200 scans/second for 60 minutes. Each session was digitally recorded. The force plate was wiped down with 70% ethanol after each session. The activity of the mouse was tracked with high accuracy, including total distance traveled, rearing events, low mobility bouts, and time spent in the center 25% of the open field. Data acquisition and calibration procedures were followed as previously described ^90^.

### Rotarod

Balance and coordination were assessed on a five-station Rotarod treadmill (IITC Life Science Inc.). Each mouse was tested three times per day for two consecutive days. All trials lasted for five minutes, the time when maximum speed was reached at a constant rate of acceleration from 4-40 rpm. A trial was terminated when a mouse fell off, made one complete backward revolution while hanging on, or after five minutes. The mice were acclimated to the room for 30 minutes on each testing day. The machine was wiped down with 70% ethanol in between each trial. Mice (10 to 14 weeks of age) were tested in four separate cohorts comprising five mice per cohort (n=10 control and n=10 *ΔSBE1/5* mutant littermates from 4 separate litters).

