## Supplementary Figures for "Combined single-cell RNA-seq profiling and enhancer editing reveals critical spatiotemporal controls over thalamic nuclei formation in the murine embryo"

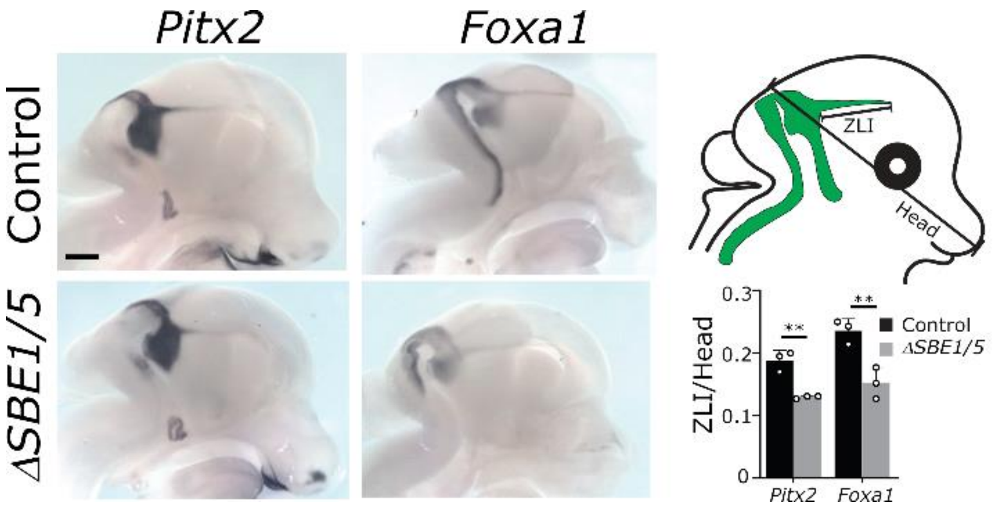


**Supplementary Figure 1. The expression of ZLI markers is maintained in *ΔSBE1/5* mutants.** Whole-mount RNA in situ hybridization for *Pitx2 and Foxa1* on bisected heads from control (top) and *ΔSBE1/5* embryos (bottom) at E12.5 showing expression in the ZLI. The area of gene expression in the ZLI, normalized to head size, is depicted at right in control and mutant embryos (**p<0.01, Student's t-test, n=3, error bars represent standard deviation). Scale bar = 500μm.

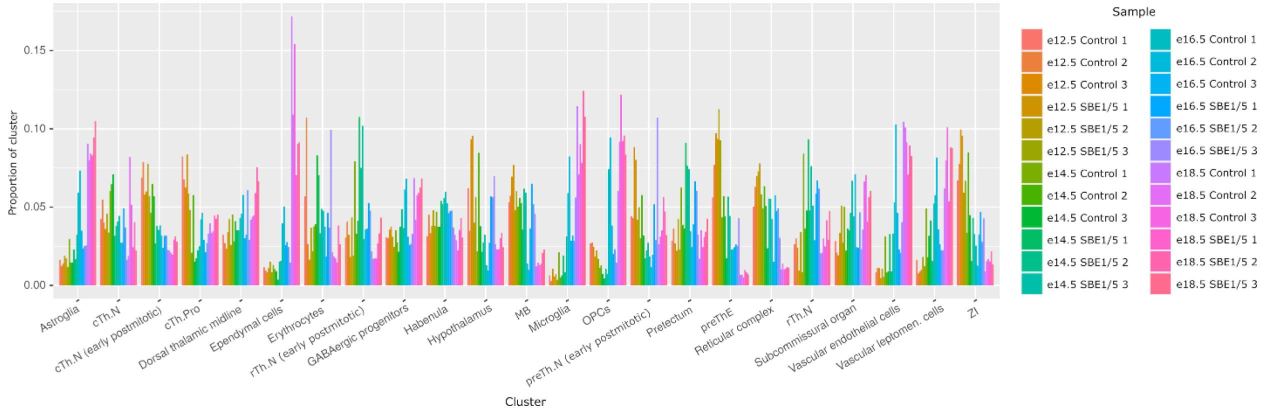


**Supplementary Figure 2. Cells from independent biological replicates are similarly delineated across all cell populations.** The barplot represents the proportion of cells from each of the 24 mice in each cluster.

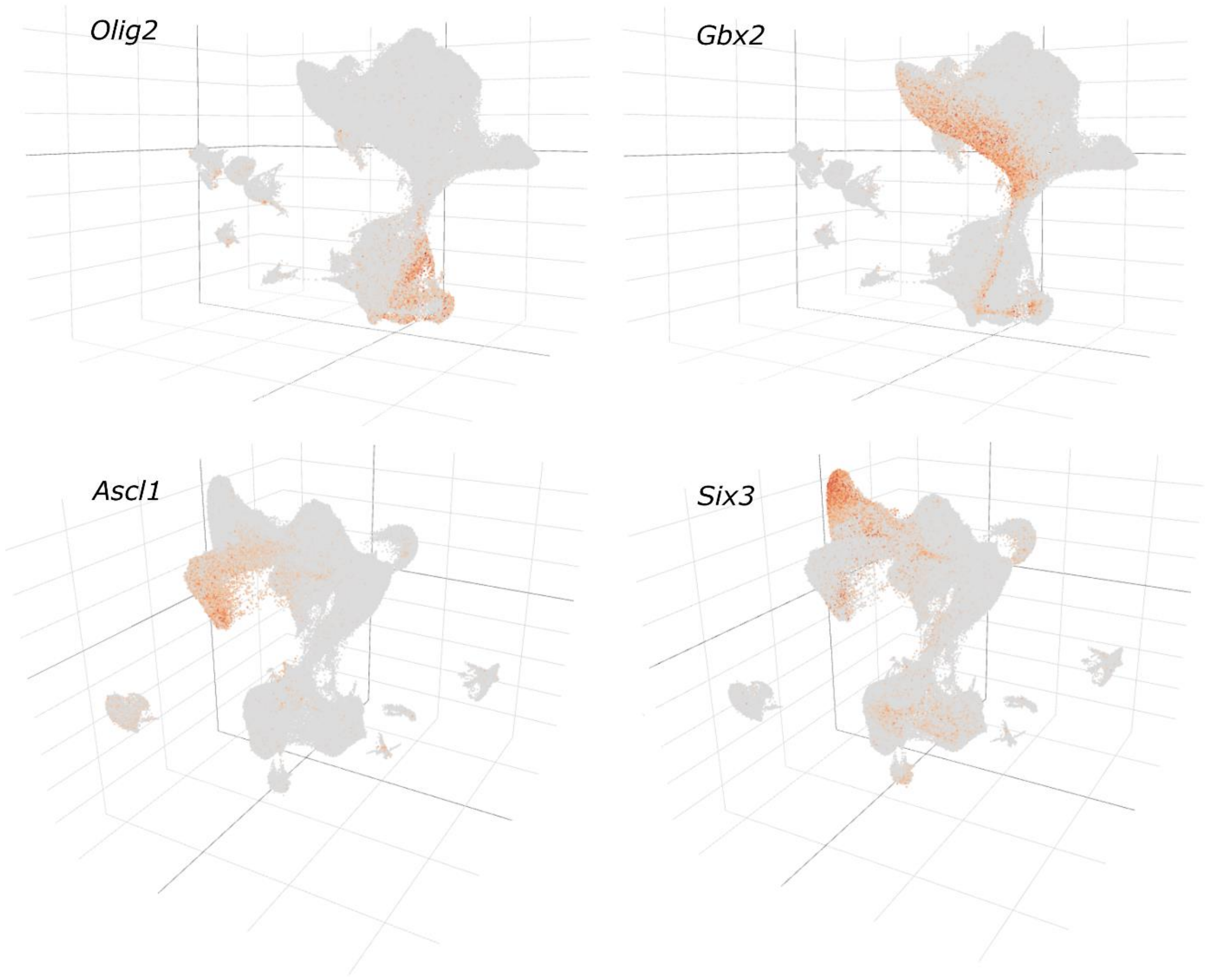


**Supplementary Figure 3. The single-cell RNA-seq atlas contains continuous developmental trajectories associated with glutamatergic and GABAergic thalamic neurons.** The 3D UMAP representation of the single-cell RNA-seq dataset is colored by the expression level of selected cTh.Pro (*Olig2*) and rTh.Pro (*Ascl1*) thalamic progenitors and cTh.N (*Gbx2*) and rTh.N (*Six3*) postmitotic thalamic neurons.

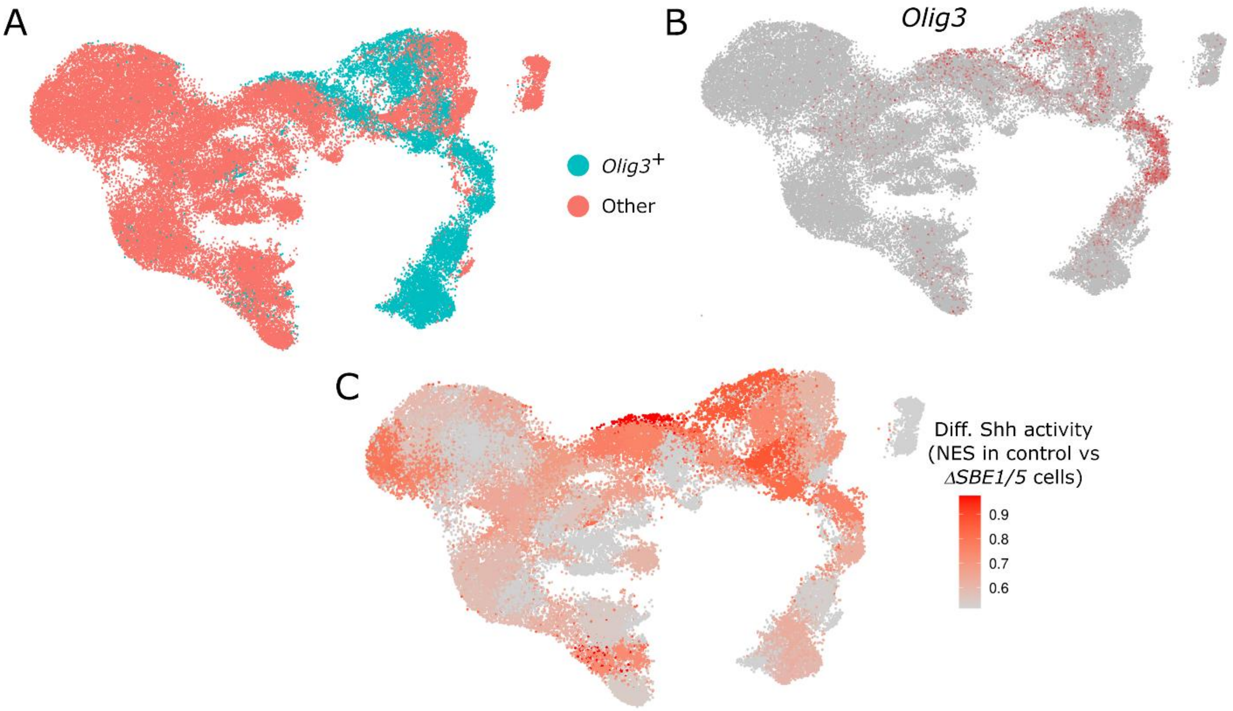


**Supplementary Figure 4. *Olig3*⁺ progenitor cell subpopulation shows a downregulation of Shh responsive genes in *ΔSBE1/5* compared to control cells.**

**A)** UMAP representation of progenitor cells from all timepoints and conditions indicating the population of *Olig3*⁺ progenitors. **B, C)** The UMAP is colored by the expression of *Olig3* (B) and the normalized enrichment score (NES) of Shh-responsive genes (*Gli1*, *Ptch1*, *Olig2*, *Nkx2-2*, *Pdlim3*, *Fst*, *Zdbf2*, *Hs3st1*, and *Slc38a11*) in control vs *ΔSBE1/5* cells (C), demonstrating the downregulation of Shh-responsive genes in *Olig3*⁺ progenitors from *ΔSBE1/5* embryos.

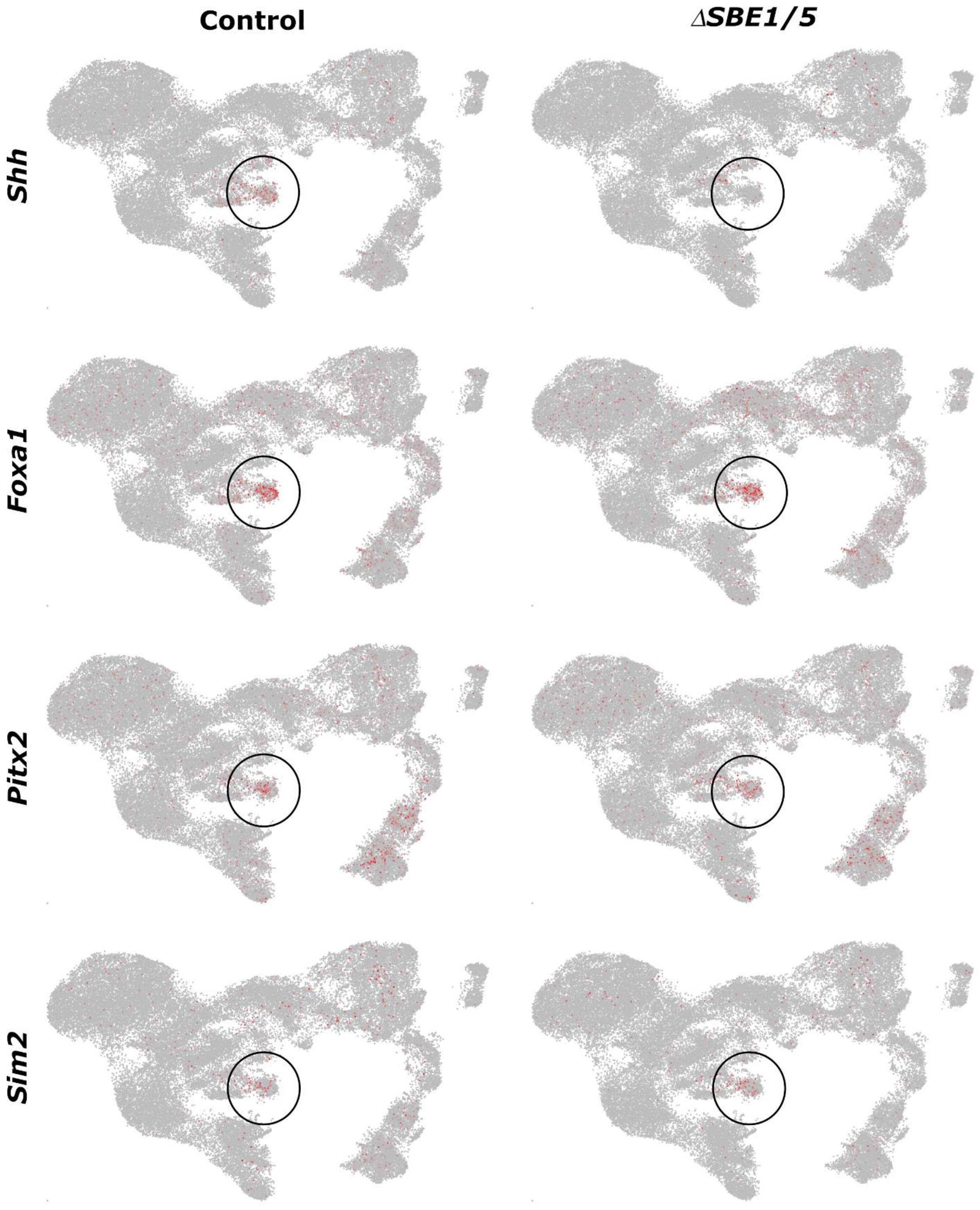


**Supplementary Figure 5. The single-cell RNA-seq data is consistent with a depletion of *Shh* expression in the ZLI of *ΔSBE1/5* mutants.** The region corresponding to the ZLI is indicated in the UMAP representation of progenitor cells from all timepoints and conditions. The UMAP is colored by the gene expression levels of *Shh* and selected transcription factors that are co-expressed in the ZLI (*Foxa1, Pitx2, Sim2*). The expression of *Shh* is downregulated in the ZLI cells of *ΔSBE1/5* embryos. However, other ZLI markers maintain their expression in *ΔSBE1/5* embryos.

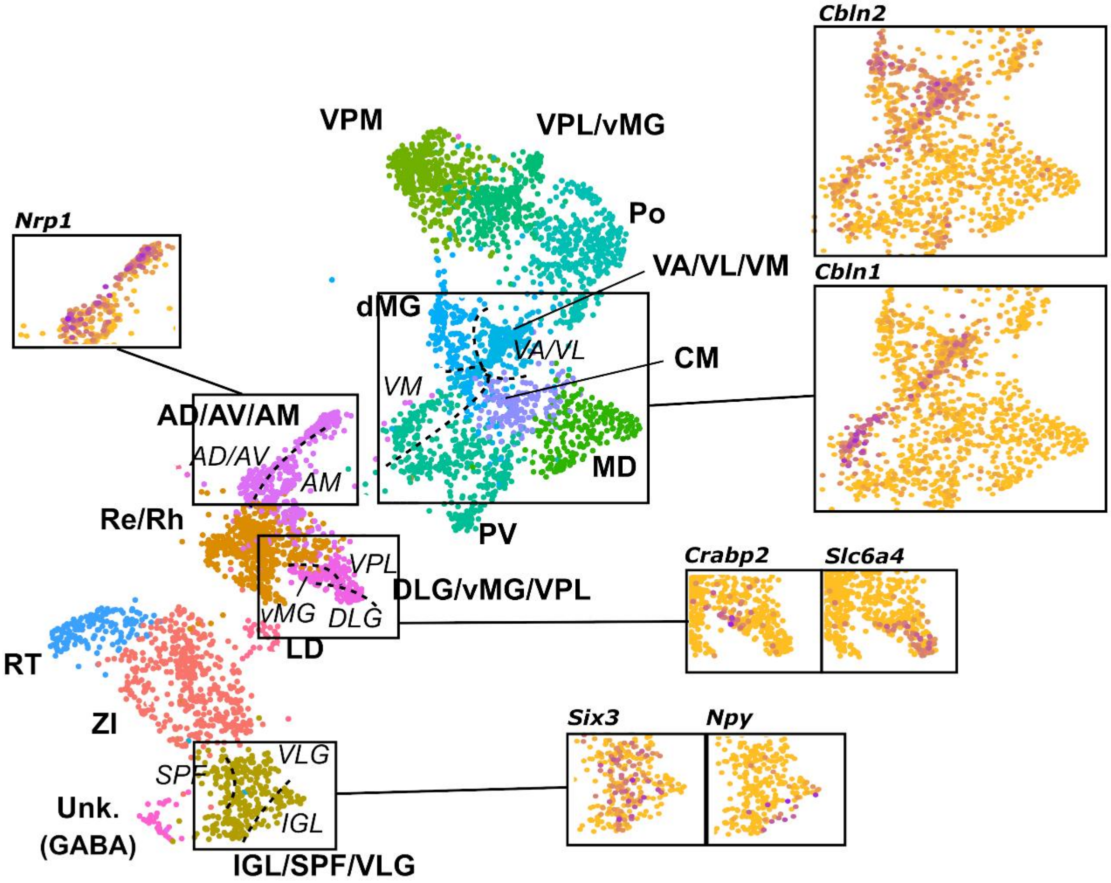


**Supplementary Figure 6. Transcriptomic signatures of closely related thalamic nuclei.** We used a spectral graph method to dissect the transcriptional heterogeneity within the cell populations identified in the clustering analysis of E18.5 thalamic nuclei. Several regions of the single-cell RNA-seq UMAP representation of E18.5 thalamic nuclei are highlighted and colored by the expression level of marker genes identified by this method.

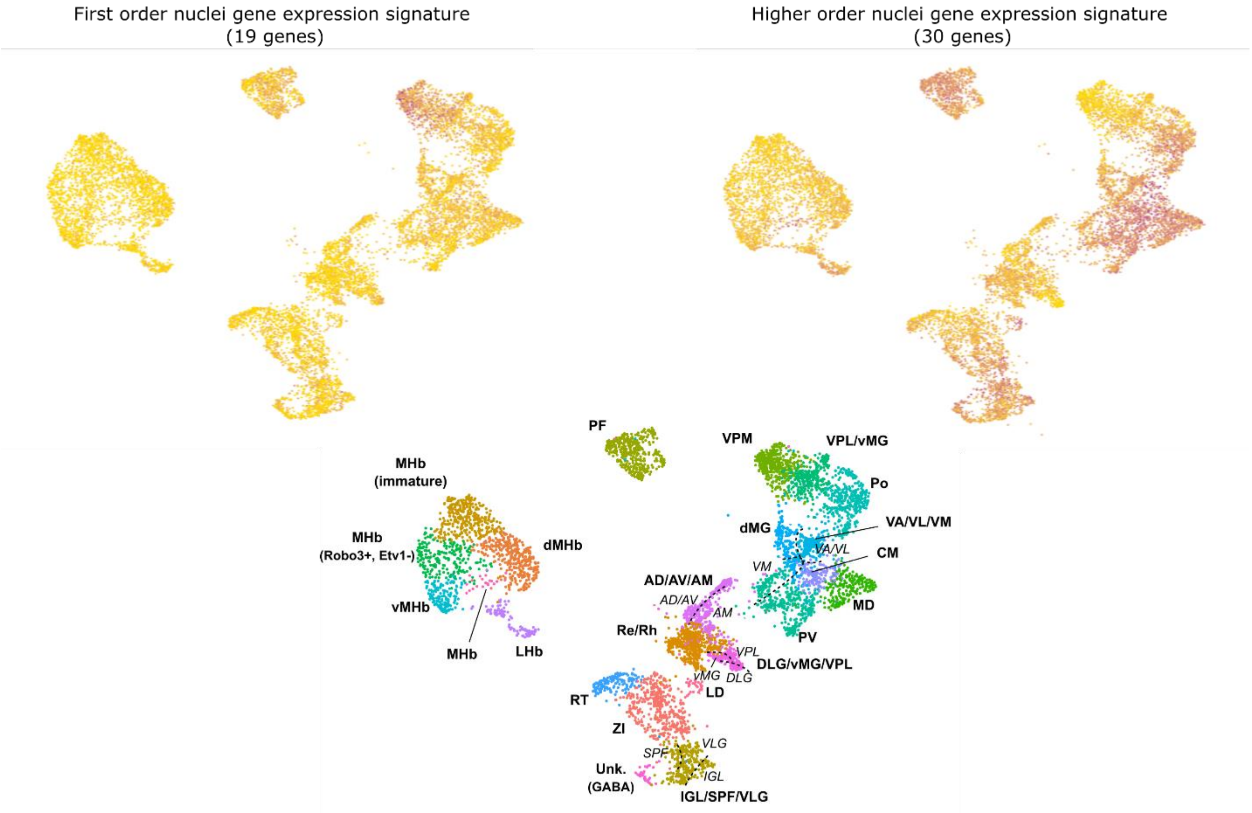


**Supplementary Figure 7. Postnatal gene expression signatures of first- and higher-order nuclei are evident at E18.5.** The single-cell RNA-seq UMAP representation of E18.5 thalamic nuclei is colored by the number of expressed genes that are postnatally associated with first order (left) and higher order (right) nuclei according to the study of Frangeul et al. 2016. For reference, the location of the different thalamic nuclei cell populations identified in this study is also indicated in the UMAP (bottom).

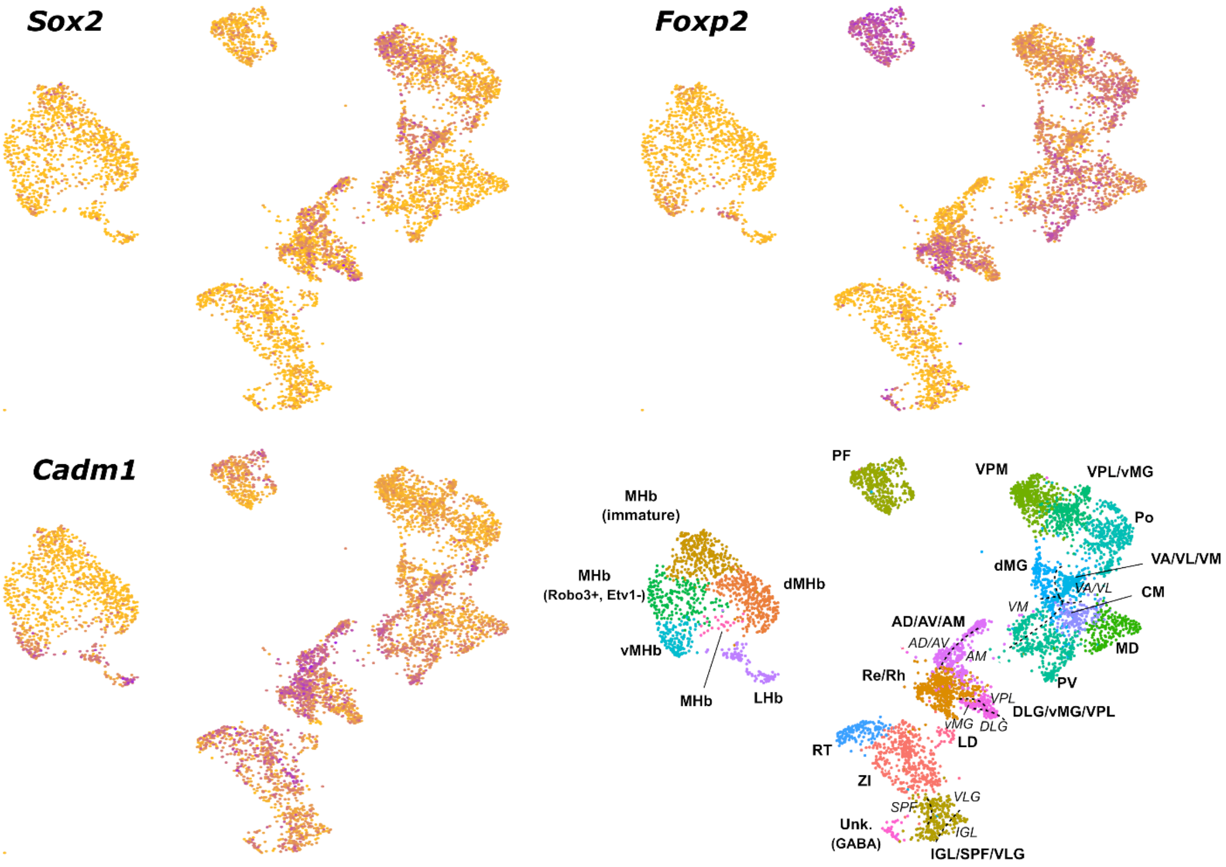


**Supplementary Figure 8. Expression of *Sox2*, *Foxp2*, and *Cadm1* lineage markers in E18.5 thalamic nuclei.** The single-cell RNA-seq UMAP representation of E18.5 thalamic nuclei cells from control mice is colored by the expression level of *Sox2*, *Foxp2*, and *Cadm1*. For reference, the location of the different thalamic nuclei cell populations identified in this study is also indicated in the UMAP (bottom).

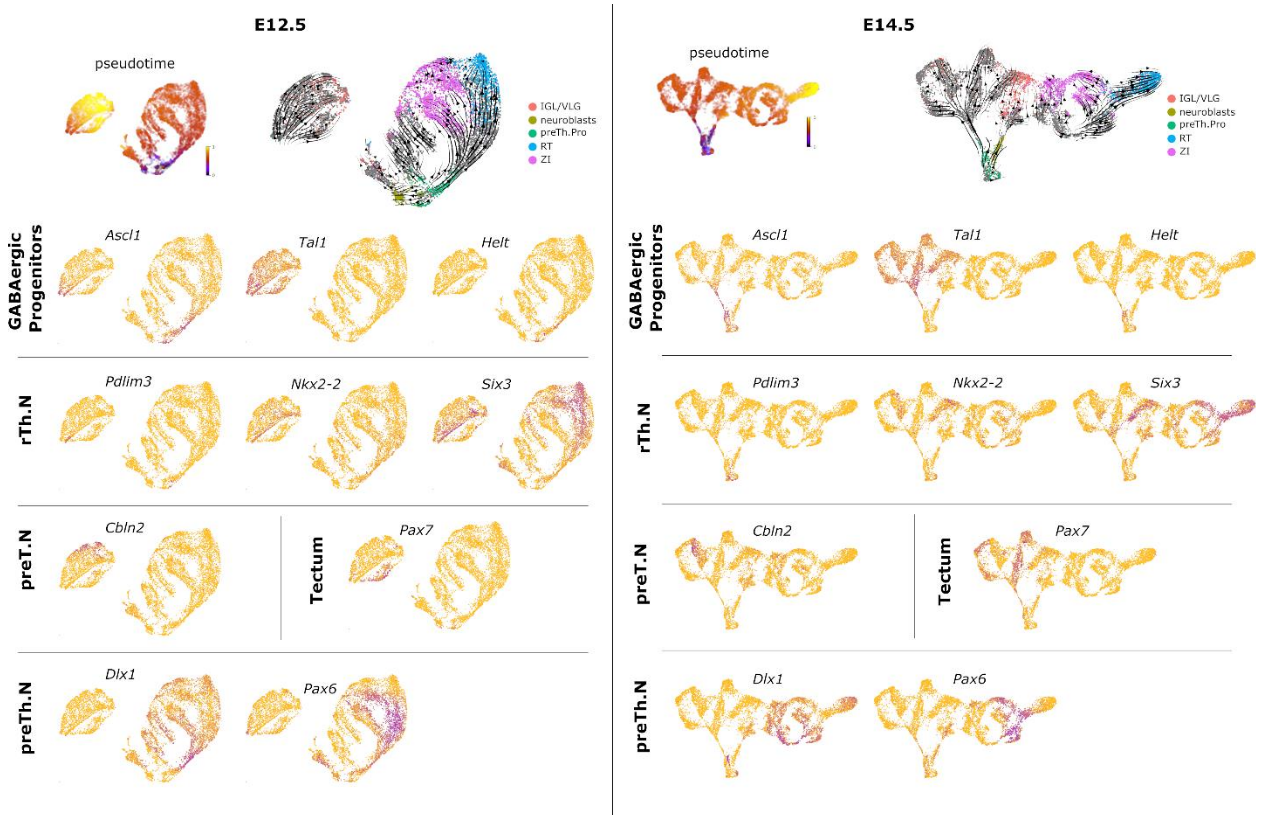


**(cont. on next page)**

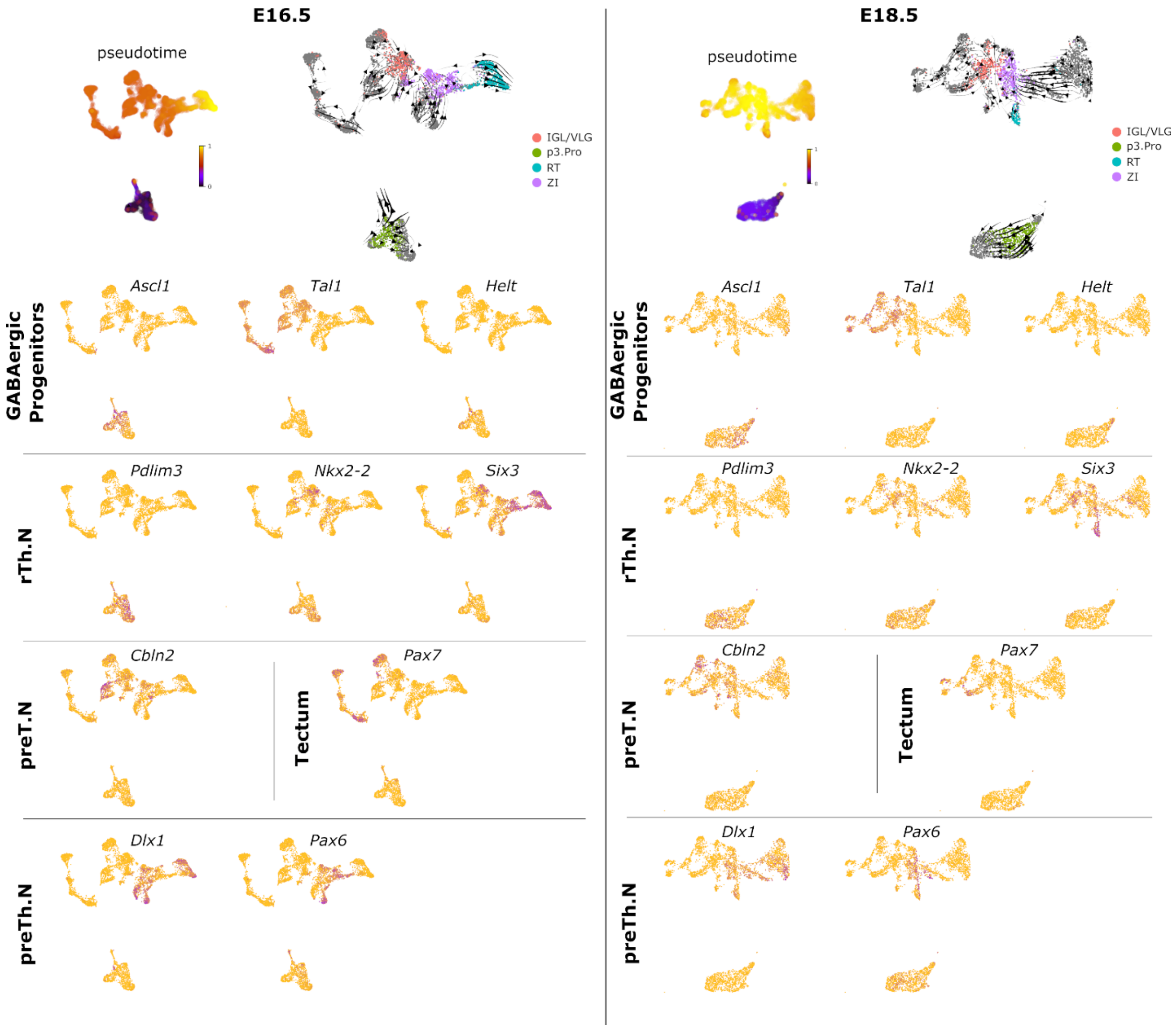


**Supplementary Figure 9. GABAergic neurons diverge from progenitors with a shared transcriptional identity.** UMAP representation and RNA velocity field of the single-cell RNA-seq data of the GABAergic cell lineages in control embryos at each developmental stage. For reference, the same UMAP is also colored by the inferred cell differentiation pseudo-time and the gene expression levels of several marker genes associated with distinct sub-lineages. The lineage originating from *Helt*$^{+}$ progenitors is characterized by the early expression of *Dlx1* followed by *Pax6* expression and gives rise to the GABAergic neuronal populations of the ZI and RT. The lineage originating from *Tal1*$^+$ progenitors splits into three distinct sub-lineages that are marked by the expression of *Six3*, *Cbln2*, and *Pax7* and give rise to GABAergic neurons of the IGL/VLG thalamic nuclei, the pretectum, and the tectum, respectively.

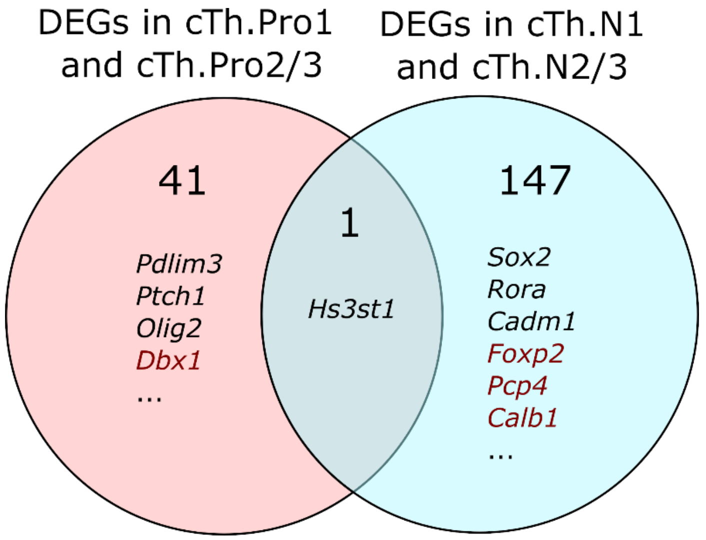


**Supplementary Figure 10. Most of the genes expressed in glutamatergic thalamic progenitor cells ceased to be expressed after mitotic arrest.** Venn diagram showing the number of genes that are differentially expressed between cTh.Pro1 and cTh.Pro2/3 progenitors (pink circle), and between cTh.N1 and cTh.N2/3 post-mitotic neurons (blue circle). Only *Hs3st1*, which marks cTh.Pro1 progenitors, continues to be differentially expressed in cTh.N1 neurons. Genes that are upregulated in cTh.Pro1 and/or cTh.N1 are marked in black, whereas genes that are upregulated in cTh.Pro2/3 or cTh.N2/3 are marked in red.

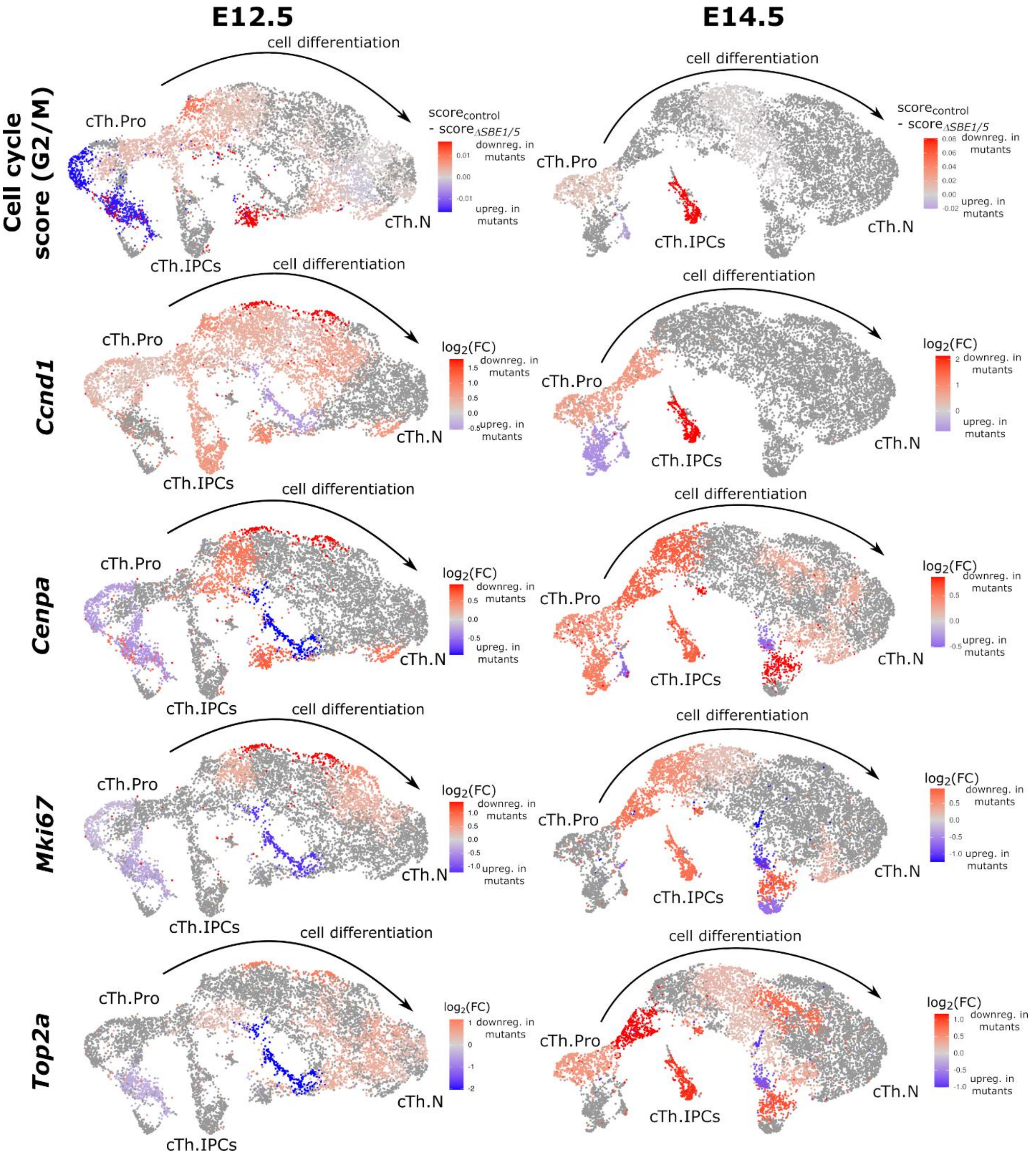


**Supplementary Figure 11. Differential gene expression analysis between control and *ΔSBE1/5* glutamatergic progenitor cells reveals the downregulation of cell cycle genes in mutant cells.** The single-cell RNA-seq UMAP representation of the glutamatergic thalamic cell lineage at E12.5 (left) and E14.5 (right) is colored by the difference in the G2/M cell cycle gene expression score between control and *ΔSBE1/5* cells (top), and the fold-change (FC) in the expression of individual cell cycle genes. The analysis shows G2/M cell cycle genes are downregulated in thalamic progenitors and IPCs in *ΔSBE1/5* embryos.

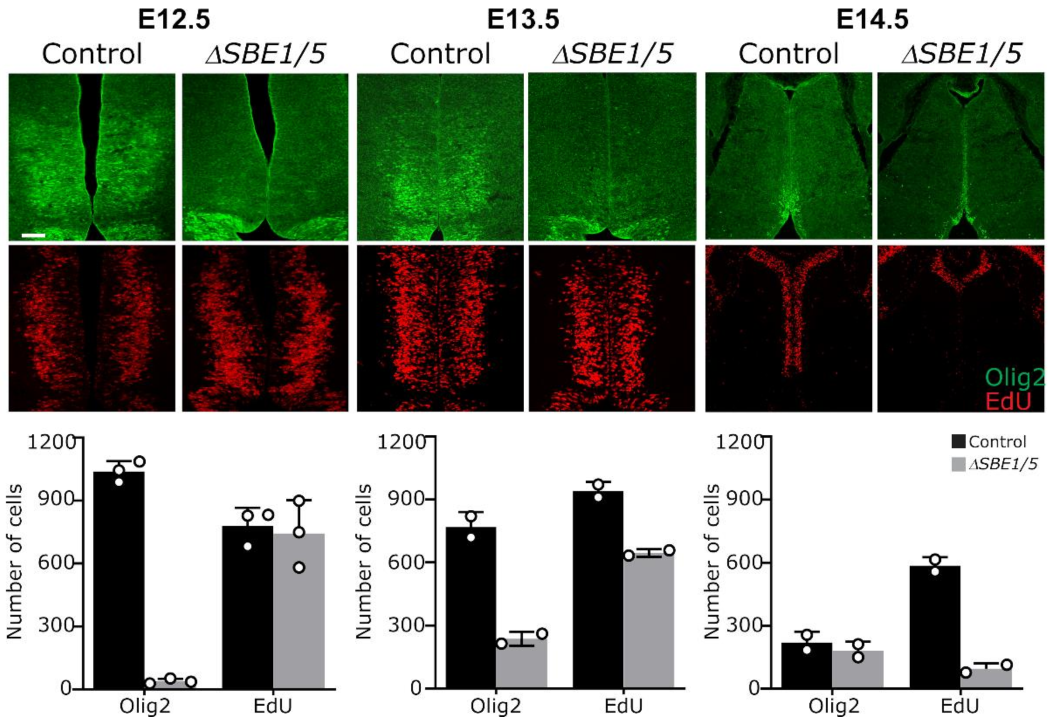


**Supplementary Figure 12. EdU incorporation and immunofluorescence staining for Olig2 in coronal sections of the thalamus in control and *ΔSBE1/5* embryos.** The figure shows a depletion of Olig2 expressing cells (cTh.Pro1 cells) and a reduction in the number of EdU incorporating cells over time in *ΔSBE1/5* embryos.

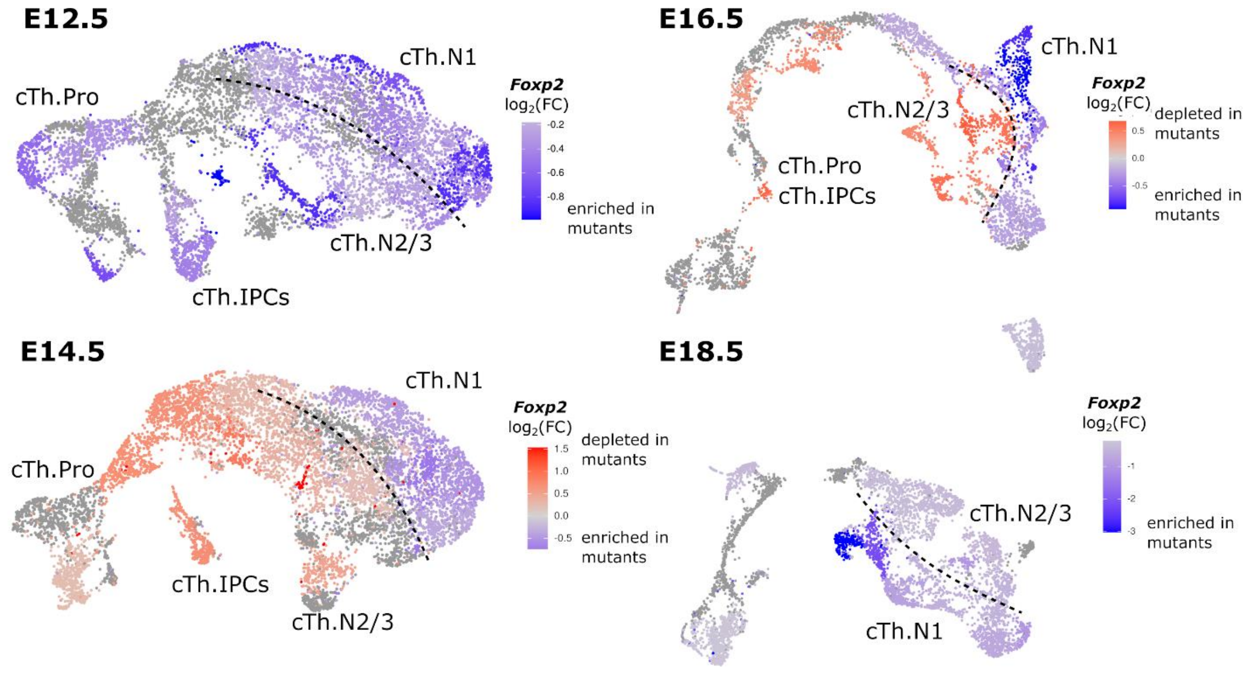


**Supplementary Figure 13. Postmitotic *Foxp2*$^+$ thalamic neurons are enriched in *ΔSBE1/5* mice.** Fold-change in the proportion of *Foxp2*$^+$ cells during glutamatergic thalamic nuclei differentiation in control and *ΔSBE1/5* mice for each timepoint, showing an enrichment for Foxp2+ cells in the cTh.N1 lineage.

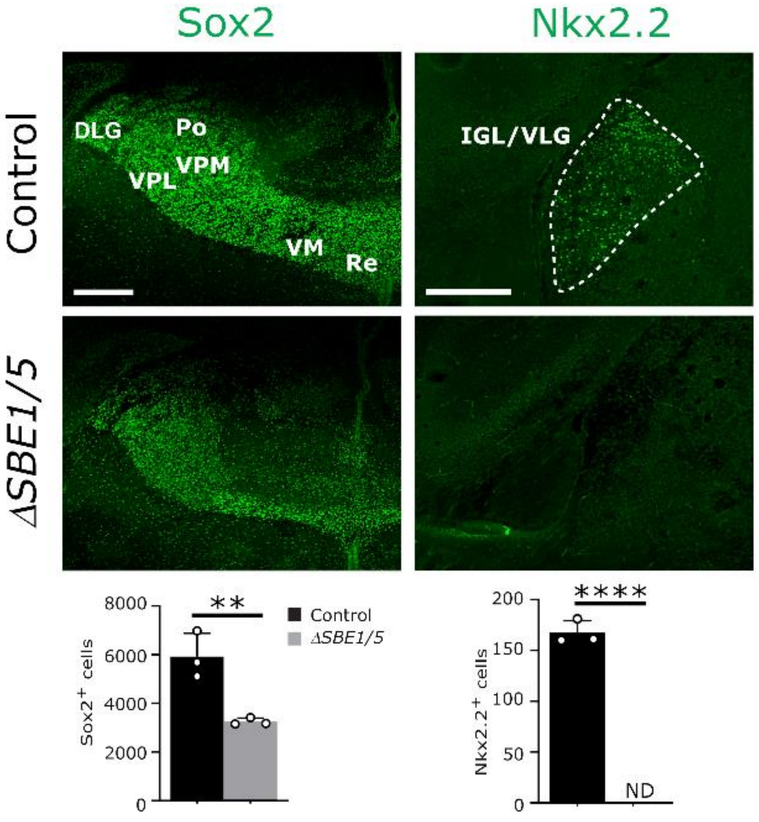


**Supplementary Figure 14. Shh-dependent nuclei are substantially reduced in newborn *ΔSBE1/5* mice.** Immunostaining for Sox2 and Nkx2.2 on coronal sections of the thalamus from control and *ΔSBE1/5* mice at P0, showing a large reduction in the size of Shh-dependent thalamic nuclei, consistent with the results of the single-cell RNA-seq analysis (**$p<0.01$, ****$p<0.0001$, Student's t-test, n=3, error bars represent standard deviation). Sox2 scale bar = 200μm; Nkx2.2 scale bare = 100μm.

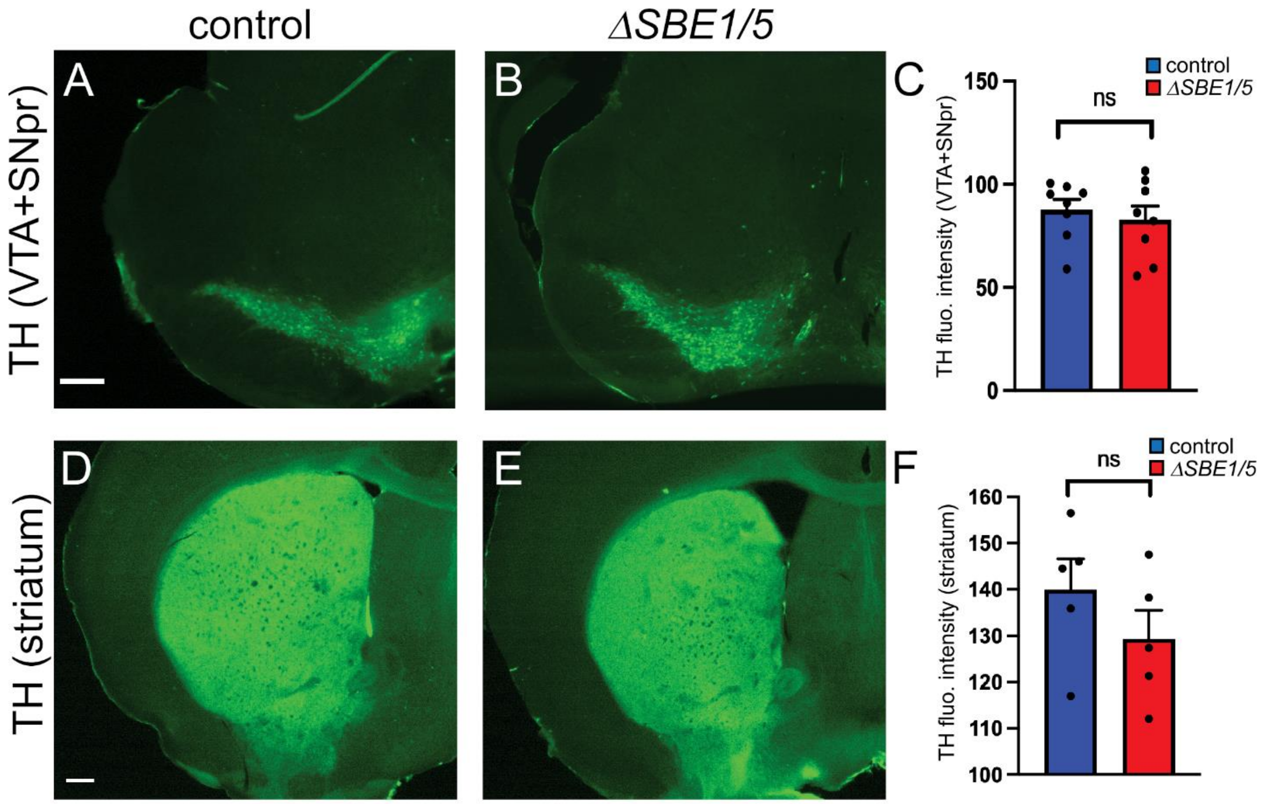


**Supplementary Figure 15. Dopaminergic neurons in the substantia nigra and their projections to the striatum are unaffected in *ΔSBE1/5* mice. (A-C)** TH immunostaining in the substantia nigra and ventral tegmental area shows no significant difference between *ΔSBE1/5* and control mice (p=0.485 n=3, 2-4 sections/brain, two-tailed student's paired t-test). **(D-F)** TH immunostaining in the striatum is unaffected in *ΔSBE1/5* compared to control mice (p=0.075 n=3, 1-2 sections/brain). Scale bars= 300 μm.

**Supplementary Tables**

**Supplementary Table 1. Differentially expressed genes between control and *ΔSBE1/5 Olig3*$^{+}$ progenitor cells.** Genes with a positive logFC have higher expression in control embryos, and genes with a negative logFC have higher expression in *ΔSBE1/5* embryos. [Provided as a separate file].

**Supplementary Table 2. Differentially expressed genes between control and *ΔSBE1/5* ZLI cells.** Genes with a positive logFC have higher expression in control embryos, and genes with a negative logFC have higher expression in *ΔSBE1/5* embryos. [Provided as a separate file].

**Supplementary Table 3. Differentially expressed genes in thalamic nuclei of E18.5 control embryos.** [Provided as a separate file].

**Supplementary Table 4. Differentially expressed genes in thalamic progenitors of E12.5 control embryos.** [Provided as a separate file].
